# Ecogenomics and functional biogeography of the *Roseobacter* group in the global oceans based on 653 MAGs and SAGs

**DOI:** 10.1101/2025.07.01.662563

**Authors:** Yanting Liu, Thorsten Brinkhoff, Meinhard Simon

**Affiliations:** Institute for Chemistry & Biology of the Marine Environment, School of Mathematics and Science, Carl von Ossietzky University Oldenburg, Carl von Ossietzky Str. 9-11, D-26129 Oldenburg, Germany; Helmholtz Institute for Functional Marine Biodiversity at the University of Oldenburg (HIFMB), Im Technologiepark 5, D-26129 Oldenburg, Germany

**Author notes:** College of Ocean and Earth Sciences, Xiamen University, Xiamen, PR China. Authors for correspondence: Yanting Liu; Meinhard Simon.

**Keywords:** *Rhodobacteraceae*, metagenome-assembled genomes, phylogenomics, bacteriochlorophyll, proteorhodopsin, genome streamlining, genome content, biogeography

## Abstract

**Background:** The *Roseobacter* group is a major component of prokaryotic communities in the global oceans. Information on this group is based predominantly on isolates and their genomic features and on the 16S rRNA gene. Assessments of prokaryotic communities in the pelagic of the global oceans indicated an unveiled diversity of this group but studies of the diversity and global biogeography of the entire group are still missing. Hence, we aimed at a comprehensive assessment of the *Roseobacter* group in the global oceans on the basis of MAGs and SAGs.

**Results:** The obtained 610 MAGs and 43 SAGs of high quality were subjected to in-depth analyses of their phylogeny, genomic and functional features. The recruitment locations range from the tropics to polar regions, include all major ocean basins. The phylogenetic analysis delineated the known RCA cluster and five pelagic clusters, two of which were completely novel: TCR (Temperate and Cold *Roseobacter*), AAPR (Arctic-Atlantic-Pacific *Roseobacter,* novel), AAR (Arctic-Atlantic *Roseobacter,* novel), COR (Central Oceanic *Roseobacter*), LUX (*Cand*. Luxescamonaceae) cluster. These clusters account for ∼70% of all *Roseobacter* MAGs and SAGs in the epipelagic. The TCR, AAPR, AAR and LUX clusters are among the most deeply branching lineages of the *Roseobacter* group. These clusters and several sublineages of the RCA and COR clusters exhibit distinct features of genome streamlining, i.e. genome sizes of <2.9 Mbp and G+C contents of <40%. The clusters exhibit differences in their functional features and also compared to other lineages of the *Roseobacter* group. Proteorhodopsin is encoded in most species of the AAPR, AAR, TCR and RCA clusters and in a few species of the COR cluster, whereas in most species of the latter, the LUX cluster and in a few species of the RCA cluster aerobic anoxygenic photosynthesis was encoded. Biogeographic assessments showed that the AAPR, AAR, TCR and RCA clusters constitute the *Roseobacter* group in the temperate to polar regions to great extent whereas the COR and LUX clusters in the tropics and subtropics.

**Conclusions:** Our comprehensive analyses shed new light on the diversification, genomic features, environmental adaptation, and global biogeography of a major lineage of pelagic bacteria.

## Background

The *Roseobacter* group is one of the major components of prokaryotic communities in the global oceans and it encompasses marine *Rhodobacteraceae* [1]. The great majority of this group was reclassified as *Roseobacteraceae* [2] but excluded its most deeply branching lineages. The abundance of the *Roseobacter* group constitutes 5-20% and in several coastal, temperate, and polar regions > 20% of the total community [3–7]. It includes members living in close association with phytoplankton algae [8–10] as well as free-living representatives such as the abundant RCA cluster [7,11] and is a major driver in the cycling of organic sulfur compounds [12]. Information on this important group of marine bacteria is based predominantly on isolates and their genomic features and on marker genes such as the 16S rRNA gene and its amplicon sequence variants (ASVs). Metagenomic and -transcriptomic studies further deepened our understanding of this group as they also captured valuable information on uncultured members and populations of members difficult to culture [4,13–16]. Assessments of pelagic prokaryotic communities based on ASVs in the Atlantic and Pacific Ocean indicate an unveiled diversity in the *Roseobacter* group as 64% of the *Rhodobacteraceae* ASVs were not affiliated to any known genus and only identified as unknown *Rhodobacteraceae* [17].

The genome sizes of the isolates span from 2.2 Mbp of *Rhodobacterales* bacterium HTCC2255 to > 5.0 Mbp of several species isolated from different marine habitats [18]. There is evidence that the prominent members of the *Roseobacter* group in pelagic prokaryotic communities have a genome size not exceeding 4.2 Mbp. Billerbeck et al. [4] assessed genomic features of prominent pelagic members, including *Rhodobacterales* bacterium HTCC2255, *Planktomarina temperata* RCA23, the type strain of the RCA cluster, and *Cand*. Planktomicrobium forsetii strain SB2, a member of the abundant CHAB-I-5 cluster. These authors found that all members of this Pelagic *Roseobacter* Cluster share a genome content and that the genome size ranges between 2.2 and 4.1 Mbp. A recent in-depth analysis of the RCA cluster, based on the analysis of metagenome assembled genomes (MAG) recruited from the global oceans, revealed that this cluster exhibits a surprising diversity including three genera and 13 species with genome sizes ranging from 1.9 to 3.1 Mbp and a distinct global biogeography of most species [7].

An evolutionary model, based on genomic features of isolates, predicts that the *Roseobacter* group evolved from a most recent common ancestor (MRCA) with other marine alphaproteobacterial lineages, including the SAR11 clade/*Pelagibacterales*, the SAR116 clade/*Puneispirillales* and *Rhodospirillales* [19]. The MRCA was predicted to have a rather large genome. During evolution, the *Roseobacter* group underwent events of gains and losses of gene families resulting in the diverse genomic features present today. Only *Rhodobacterales* bacterium HTCC2255 escaped major events of gene gains, and losses of gene families prevailed, yielding its small genome which exhibits features of genome streamlining. In contrast to the *Roseobacter* group, the other orders derived from this MRCA are characterized predominantly by gene losses such that their genome sizes continuously decreased to the present size range of 1.3 to 3.1 Mbp [18,20,21]. Typical features of their small and streamlined genomes include a low G+C content (30-40%), a high coding density (> 91%), and a small proportion of pseudogenes (< 10% of total genes) [20–22]. Similar features of genome streamlining were reported from MAGs and SAGs of the *Ca*. Luxescamonaceae family, encoding genes of aerobic anoxygenic photosynthesis (AAP) and carbon fixation via the Calvin-Benson-Bassham (CBB) cycle [23,24]. The authors proposed it as a sister family to *Rhodobacteraceae*.

Features of genome streamlining have been documented in the *Roseobacter* group in the genomes of *Rhodobacterales* bacterium HTCC2255 and the RCA cluster (see above). There is evidence that genome streamlining also occurred in other so far little or unknown lineages of this group as the G+C content of single amplified genomes (SAG) and metagenome assembled genes binned to the *Roseobacter* group exhibit G+C contents of < 40% [13,18,19,22]. As the RCA cluster and these SAGs branch off from intermediate regions of the phylogenomic tree of the *Roseobacter* group streamlining events to adapt to the energy- and nutrient-limited near surface ocean appear to have happened at least twice. Further, there is evidence from the RCA cluster that these adaptations to specific environmental conditions lead to distinct global biogeographic patterns of sublineages [7].

Hence the aim of this study was a comprehensive assessment of the *Roseobacter* group in the global oceans on the basis of MAGs and SAGs recruited from databases of the Ocean Microbiomics (https://microbiomics.io/ocean/), the COGITO project (PRJEB28156), and an Atlantic Ocean transect [25]. The main questions we asked was whether the *Roseobacter* group encompasses so far unknown sublineages and whether the diversity of known pelagic clusters is already captured. As we identified four novel sublineages/clusters and multiple novel genomospecies (termed species herein) we further aimed i) to assess their genomic features and within cluster diversification, ii) to elucidate their functional features and iii) to assess the global biogeography of the different cluster members. As a result, we identified two novel clusters and a greatly enhanced diversification of three other clusters: The most deeply branching cluster with seven species and including *Rhodobacterales* bacterium HTCC2255, two completely new deeply branching clusters without any known taxonomic signature, and another large cluster consisting of six subclusters and 19 species including isolates *Cand*. Planktomicrobium forsetii strain SB2 and HIMB11 as members. Within each cluster sublineages exhibit features of genome streamlining and include members encoding proteorhodopsin (PR) for complementary light energy acquisition. Cluster members exhibit distinct global biogeographies reaching out into tropical and subtropical regions in which the *Roseobacter* group has not been detected previously.

These analyses complement comparable studies on other major prokaryotic orders and families in the epipelagic ocean like *Prochlorococaceae* [26–28] *Pelagibacterales*/SAR11 [20,29,30] the SAR116 clade/*Puneispirillales* [21], the SAR86 clade [31] and the SAR202 clade [32].

## Materials and methods

### Sample source, metagenomic quality control, assembly and binning

We collected 133 *Roseobacter* SAGs and 1290 MAGs from the Ocean Microbiomics Database, as well as 40 MAGs from the North Sea (COGITO project, PRJEB28156), based on the genome taxonomy database (GTDB, r207) (Supplementary Table S1). Additionally, 40 MAGs were reconstructed from 22 metagenomic samples of an Atlantic Ocean transect from 62°S to 47°N [25] (Supplementary Table S1). Metagenomic binning and taxonomic classification of these MAGs followed methods described previously [7]. Genome quality was assessed for completeness and contamination using CheckM (version 1.0.13) [33] and Anvi’o (version 5.5.0) [34]. Forty-three SAGs and 610 MAGs passed the quality filter of a mean completeness of ≥ 70%, a mean contamination of ≤ 5% and scaffolds with an N50 of ≥ 10 kb and were selected for downstream analyses (Supplementary Table S2).

### Phylogenetic analyses

#### Genome-based phylogeny of the Roseobacter group

We carried out a phylogenomic analysis of 154 representative genomes of our dataset after using dRrep (version 3.4.3) [35] and 344 reference genomes of the *Roseobacter* group and added two outgroup genomes (*Agrobacterium tumefaciens* WRT31 and *Agrobacterium fabacearum* P4). The phylogenetic tree was inferred using 120 conserved bacterial marker genes. The identification and alignment of marker genes and trimming of a concatenated alignment were performed following the GTDB-Tk workflow (version 2.1.1) [36]. The resulting concatenated alignment was applied to construct a maximum-likelihood phylogenetic tree using IQ-TREE (version 2.3.4) [37] under the best-fit model Q.yeast+F+I+R10 substitution model. Tree robustness was evaluated with 1000 ultrafast bootstrap replicates. The final tree was visualized and annotated using the Interactive Tree of Life view (iTOL) [38].

#### Phylogenetic tree construction of the proteorhodopsin and pufM genes

For rhodopsin gene analysis, 526 xanthorhodopsin (XR) and 367 PR sequences were downloaded from the UniProt database (https://www.uniprot.org/). Sequences shorter than 200 amino acids were excluded and remaining sequences were clustered at 90% amino acid identity using MMseqs2 (version 14.7e284) [39], resulting in 214 PR and 300 XR representative reference sequences. Additionally, 207 PR gene sequences were extracted from the *Roseobacter* group genomes analyzed in this study. These sequences, along with the references, were aligned with MAFFT (version 7.407) [40] with the "--auto" parameter and trimmed with trimAl (version 1.5) [41] using the "-gappyout" option to remove poorly aligned regions. A maximum likelihood phylogenetic tree was constructed using IQ-TREE [37] under the best-fit model LG+F+R8 model, with 1000 ultrafast bootstrap replicates for support.

A similar workflow was employed for the *puf*M gene phylogeny. This analysis included 448 *puf*M gene sequences, consisting of 348 reference sequences from the NCBI RefSeq database and 100 sequences retrieved from the *Roseobacter* group genomes in our dataset. The sequences were aligned with MAFFT [40], trimmed with trimAl ("-gappyout") [41] and used to infer a maximum likelihood phylogenetic tree in IQ-TREE [37] with the best-fit model LG+F+R9 model, supported by 1000 ultrafast bootstrap replicates. Both rhodopsin and *puf*M phylogenetic trees were visualized using iTOL [38].

### Analysis of genome and gene features

To estimate the genome size, G+C content, and coding density of MAGs and SAGs, we used CheckM (version 1.0.13) [33]. The genome size was further corrected based on the contamination and completeness [42]. Open reading frames (ORFs) and gene functions were predicted with Prokka (version 1.14.6) [43], while pseudogenes were identified using Pseudofinder (version 1.1.0) for each genome [44]. To identify genes associated with metabolic modules, pathways or operons, HMMER was employed to search against KOfamScan HMMs profiles [45]. The completeness of a metabolic module, pathway, or operon was assessed using KEGG-Decoder (version 1.3) [23] and METABOLIC (version 4.0) [46]. A module, pathway, or operon was considered as ‘present’ in a genome if its completeness score exceeded 75%.

Membrane transporter genes encoded within the MAGs and SAGs were identified by searching against the Transporter Classification Database (TCDB) using DIAMOND [47] with an e-value threshold of 1e -30 [48,49]. Transporter genes were categorized based on TCDB classifications: those annotated as 3.A.1 were assigned to ATP-binding cassette (ABC) transporters, 2.C.1 to TonB-dependent transporters (TBDT), 2.A.56.1 to tripartite ATP-independent periplasmic (TRAP) transporters, and 1.A.80.1 to tripartite tricarboxylate transporters (TTT). Substrate specificity of ABC transporters was further checked with KEGG annotations using the KEGG Orthology (KO) database. A transporter was considered as ‘present’ in a genome if its completeness score was higher than 80%.

### Biogeography of the *Roseobacter* group in the global ocean

To explore the global distribution of the *Roseobacter* group, 745 non-redundant shotgun metagenomic samples were collected from the Ocean Microbiomics Database [50], the OceanDNA dataset [51], the Malaspina 2010 Circumnavigation Expedition [52], an Atlantic Ocean transect from 62°S to 47°N [25] and a cruise to the Arctic and Southern Ocean [53]. These samples consisted of 518 samples from the epipelagic (< 200 m), 150 from the mesopelagic (200-1000 m) and 77 samples from the bathypelagic (> 1000 m). Each metagenomic sample was subsampled to 10 million reads (10^7^) using Seqtk (version 1.4, https://github.com/lh3/seqtk) to normalize sequencing depth. The abundance and taxonomic classification were estimated using Kraken2 (version 2.1.3) [54] and Bracken (version 2.9) [55] with default parameters. To align classifications with the GTDB, a custom GTDB-r207 database was constructed for Kraken2 using Struo2 (version 2.1) [56], enabling precise taxonomic profiling of the *Roseobacter* group across different oceanic regions.

The abundance of the genus *Amylibacter* was used to represent the abundance of the TCR cluster, as genomes from this cluster are classified within the *Amylibacter* according to the current GTDB taxonomy.

## Results and Discussion

### Phylogenetic features and recruitment regions of *Roseobacter* MAGs and SAGs

The 610 MAGs and 43 SAGs which passed the quality control were subjected to in-depth analyses of their phylogeny, genomic, and functional features. The locations from where the MAGs and SAGs were recruited span a wide range of latitudes and temperature zones ((sub)tropical, temperate, (sub)polar), ocean regions (Fig. 1A) and the epi- to the bathypelagic (Supplementary Table S2). The majority of MAGs and SAGs was recruited from the epipelagic (84%) and the Arctic and Atlantic Oceans (76%).

**Figure 1.**
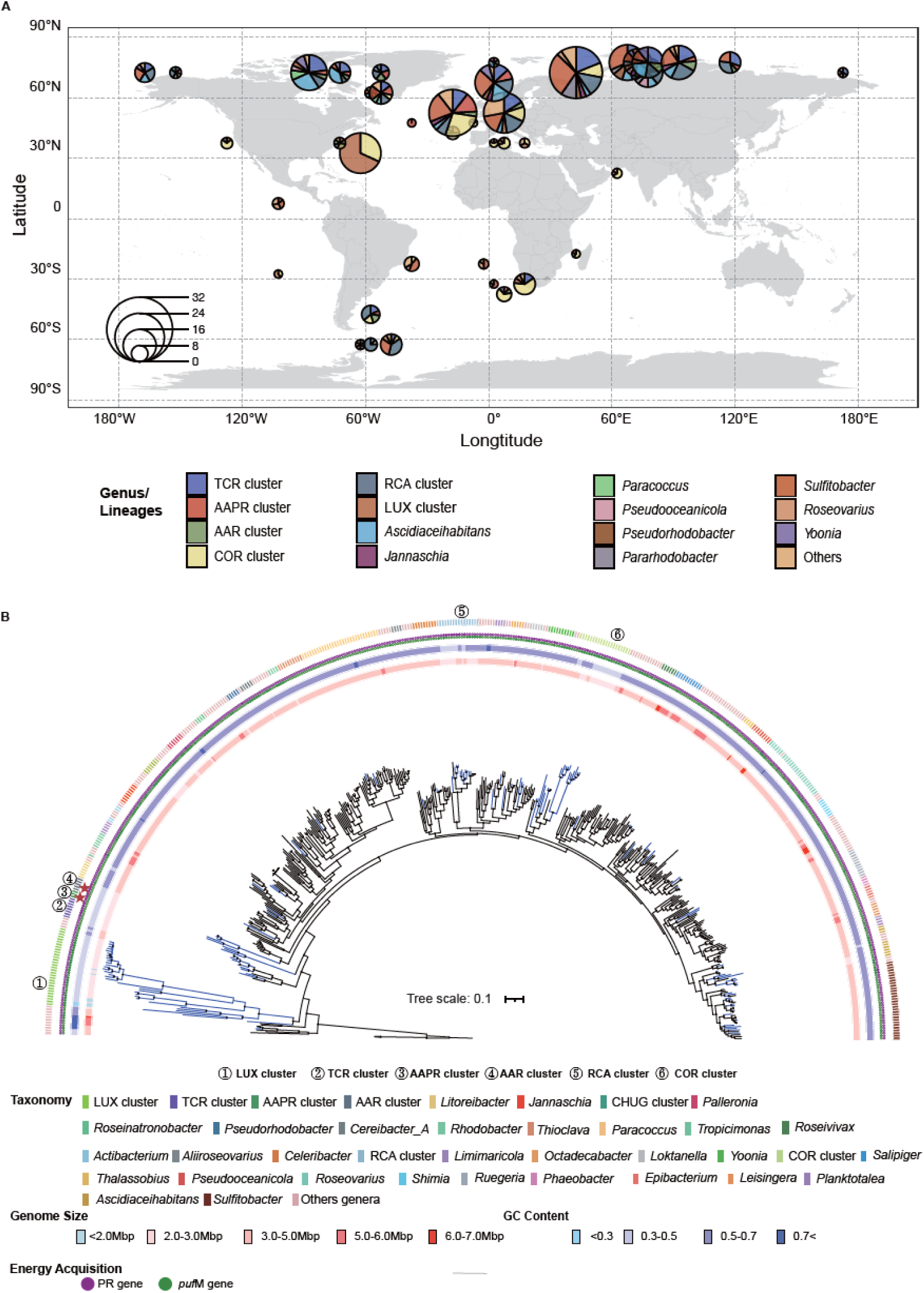
Global recruitment sites of *Roseobacter* MAGs and SAGs and phylogenetic tree of the *Roseobacter* group. (A) sites of the recruited 653 high-quality genomes. The center of each circle represents genomes recovered within the ± 5° latitude/longitude range (for exact coordinates of the locations see Table S2). The radius of the circle indicates the number of genomes. The color code refers to genomes affiliated to the pelagic clusters and other genera of the *Roseobacter* group. "Others” include the genera with less than five genomes. (B) Phylogenetic tree of the *Roseobacter* group including 498 genomes. 154 genomes are representative MAGs and SAGs recruited in this study (blue), 344 genomes are references to representative species of the *Roseobacter* group. and two are outgroup genomes (*Agrobacterium tumefaciens* WRT31, *Agrobacterium fabacearum* P4). From inner to outer semicircles: Genome size, G+C content, presence/absence of genes encoding *puf*M and PR, the genus-level classification. The pelagic clusters are labeled by numbers 1 to 6 and the two completely novel lineages (AAPR and AAR clusters) by a red star.

The phylogenetic analysis revealed two completely novel clusters and, in addition to the well-known RCA cluster, delineated five pelagic clusters based on the phylogeny and regions where their MAGs and SAGs were recruited. We propose the following names for these clusters: TCR (Temperate and Cold *Roseobacter*), AAPR (Arctic-Atlantic-Pacific *Roseobacter*), AAR (Arctic-Atlantic *Roseobacter*), COR (Central Oceanic *Roseobacter*) cluster, and LUX (*Cand*. Luxescamonaceae). These clusters account for 70% and 21% of all *Roseobacter* MAGs and SAGs in the epi- and mesopelagic, respectively. In the bathypelagic, only MAGs affiliated to *Sulfitobacter* and *Marinovum* were recruited.

The TCR, AAPR, and AAR clusters are among the most deeply branching lineages of the *Roseobacter* group, branching off near even more deeply rooted genera including *Pontivivens*, *Neptunicoccus*, *Amylibacter* and the *Cand*. Luxescamonaceae family (Fig. 1B, Supplementary Fig. S1). The TCR cluster is rooted more deeply than the two other clusters. Further, our analysis revealed that the phylogenetic position of the *Cand*. Luxescamonaceae family is within the *Roseobacter* group (Fig. 1B, Supplementary Fig. S1) and thus is another deeply branching cluster of this group. Therefore, we included it in our analyses and termed it LUX cluster. The RCA and COR clusters diverge in intermediate regions of the phylogenetic tree (Fig. 1B).

The *Roseobacter* group including our MAGs and SAGs exhibits a large range in genome size (1.45-6.33 Mbp) and G+C content (28-73%). The dereplicated 154 MAGs and SAGs have a smaller mean genome size (3.23 ± 0.93 Mbp) and lower mean G+C content (50 ± 11%) than the reference genomes of the cultured strains (mean genome size: 4.12 ± 0.70 Mbp; mean G+C content: 62 ± 5%) (Supplementary Fig. S2).

### Genomic features, phylogenetic diversity and recruitment regions of the clusters

The AAR and AAPR clusters are completely novel lineages without any previous reference. The TCR cluster includes *Rhodobacterales* bacterium HTCC2255 of which only the genome is known as the isolate got lost. The LUX cluster, proposed as a sister family to *Rhodobacteraceae* (see above) [23], is newly affiliated to the *Roseobacter* group. The RCA cluster is well known and its phylogenomic and functional diversity have been recently described based on isolates and MAGs [7]. The COR cluster includes two isolates with fully sequenced genomes (HIMB11, SB2) [4,57]. Two more isolates almost identical to SB2 have been recently described as *Cand*. Thalassiovivens spotiae [58]. These six pelagic *Roseobacter* clusters overlap partially with previously described clusters whose analyses were based on the genomes of isolates [4].

The MAG- and SAG-based clusters of the present study exhibit distinct cluster-specific genomic features (Fig. 2B, Supplementary Tables S4, S6, S8, S10, S11, S13). The AAR, AAPR and LUX clusters are characterized by small genomes (< 2.7 Mbp), < 2300 coding gene sequences (CDS) for all gene features and < 2100 CDS for intact genes, excluding pseudogenes, a low G+C content (< 39%) and a high coding density (> 90% for all gene features, > 84% for intact genes, excluding pseudogenes). The TCR, RCA, and COR clusters exhibit more variable genomic features, i.e. also larger genomes, higher CDS counts, higher G+C contents, and lower coding densities.

**Figure 2.**
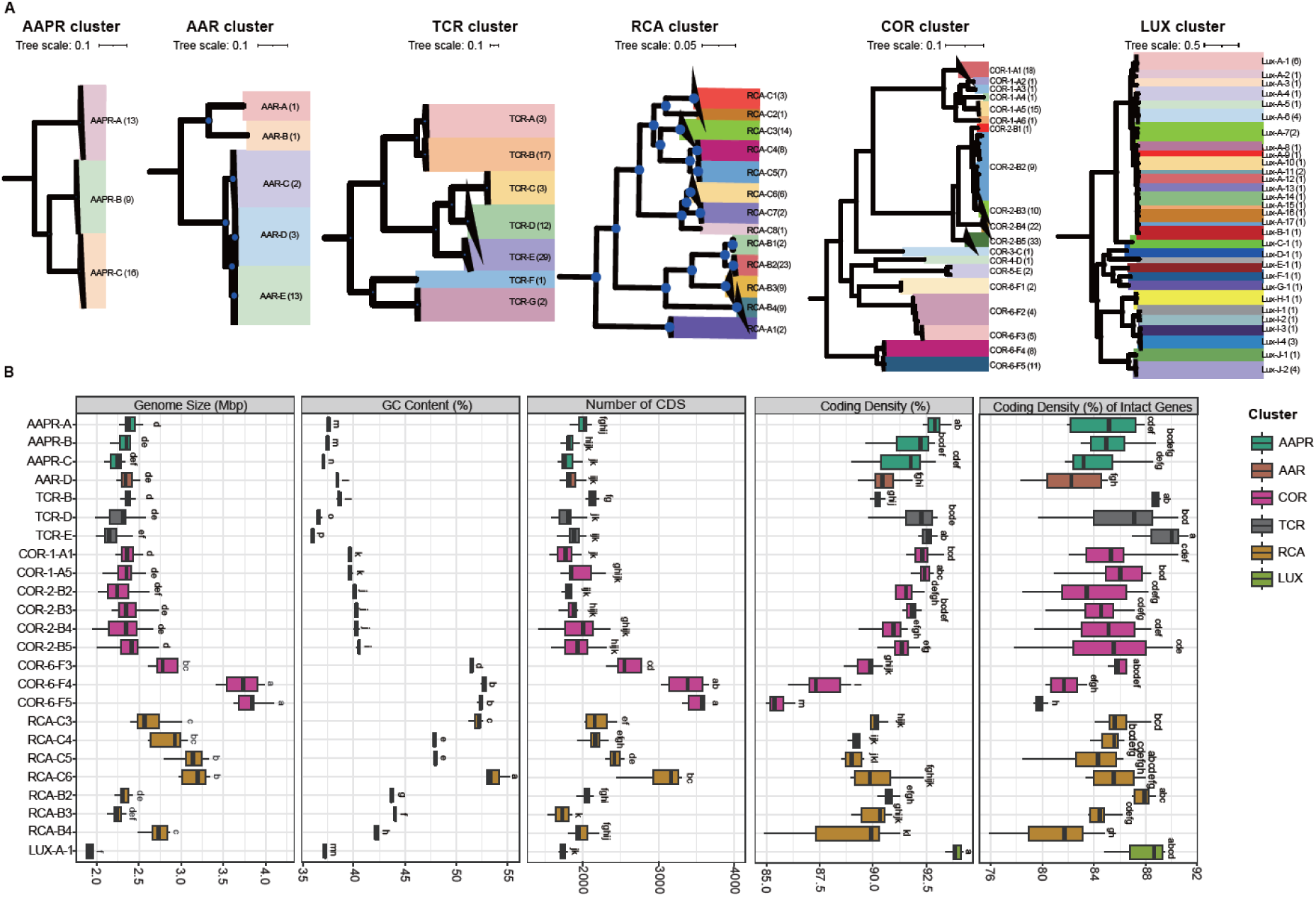
Phylogenetic tree and genomic features of six pelagic clusters of the *Roseobacter* group. (A) Phylogenetic tree of the AAPR, AAR, TCR, RCA, COR, and LUX clusters. The number of genomes recruited for each species is indicated in parenthesis. (B) Box-Whisker plots with median, 25% and 75% percentiles and range of genomic features of each species, including genome size, G+C content, number of coding sequences (CDS), coding density of total and intact genes (%). Only species with more than four genomes are included in this analysis and shown. Significant differences (p < 0.05, Tukey test) among different species are denoted by letters a to l.

### TCR cluster

The TCR cluster comprises 65 MAGs, one SAG, and the genome of *Rhodobacterales* bacterium HTCC2255. We identified seven species within this cluster, based on ANI dissimilarities of < 95% (Fig. 2A, Table 1, Supplementary Fig. S3, Supplementary Tables S4 and S5). The SAG and *Rhodobacterales* bacterium HTCC2255 affiliate with species TCR-D. Genome sizes range from 1.98 to 2.65 Mbp, CDS counts are ≤ 2213, the coding density of all gene features ranges from 88.51 to 93.03% and of intact genes from 79.73 to 91.25% (Table 1, Supplementary Table S4). Five of the seven species exhibit G+C contents < 39% whereas the two most deeply branching species, TCR-F and -G, have G+C contents of 48.29% and 50.17% and CDS counts of 2713 and 2520, respectively (Fig. 2A, Table 1). Hence, the TCR cluster shows a clear separation of five of the seven species with features typical of streamlined genomes. Statistical analyses revealed several significantly different genomic features among the different species even though only species B, D, and E could be included in these analyses (Fig. 2B).

**Table 1.**
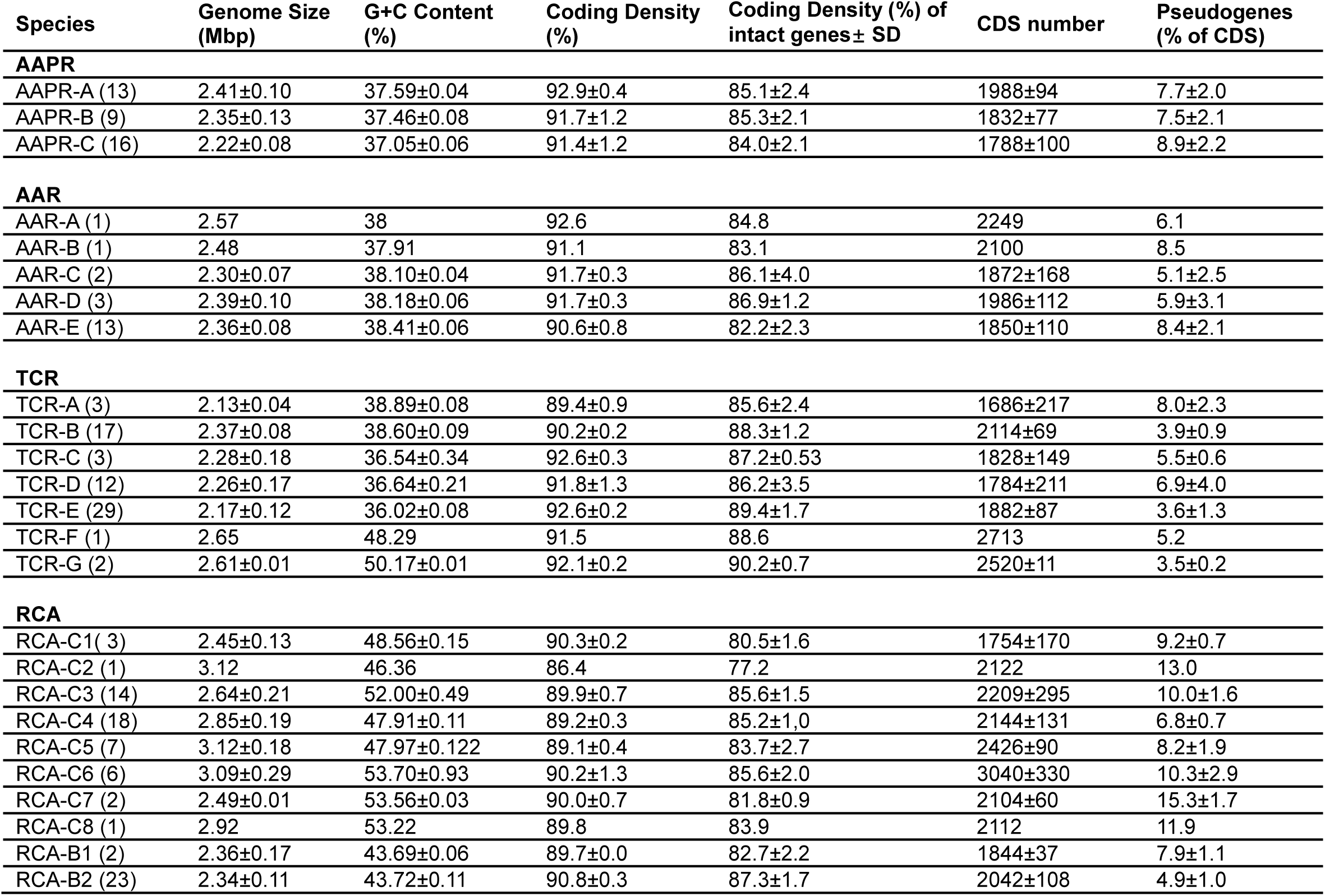
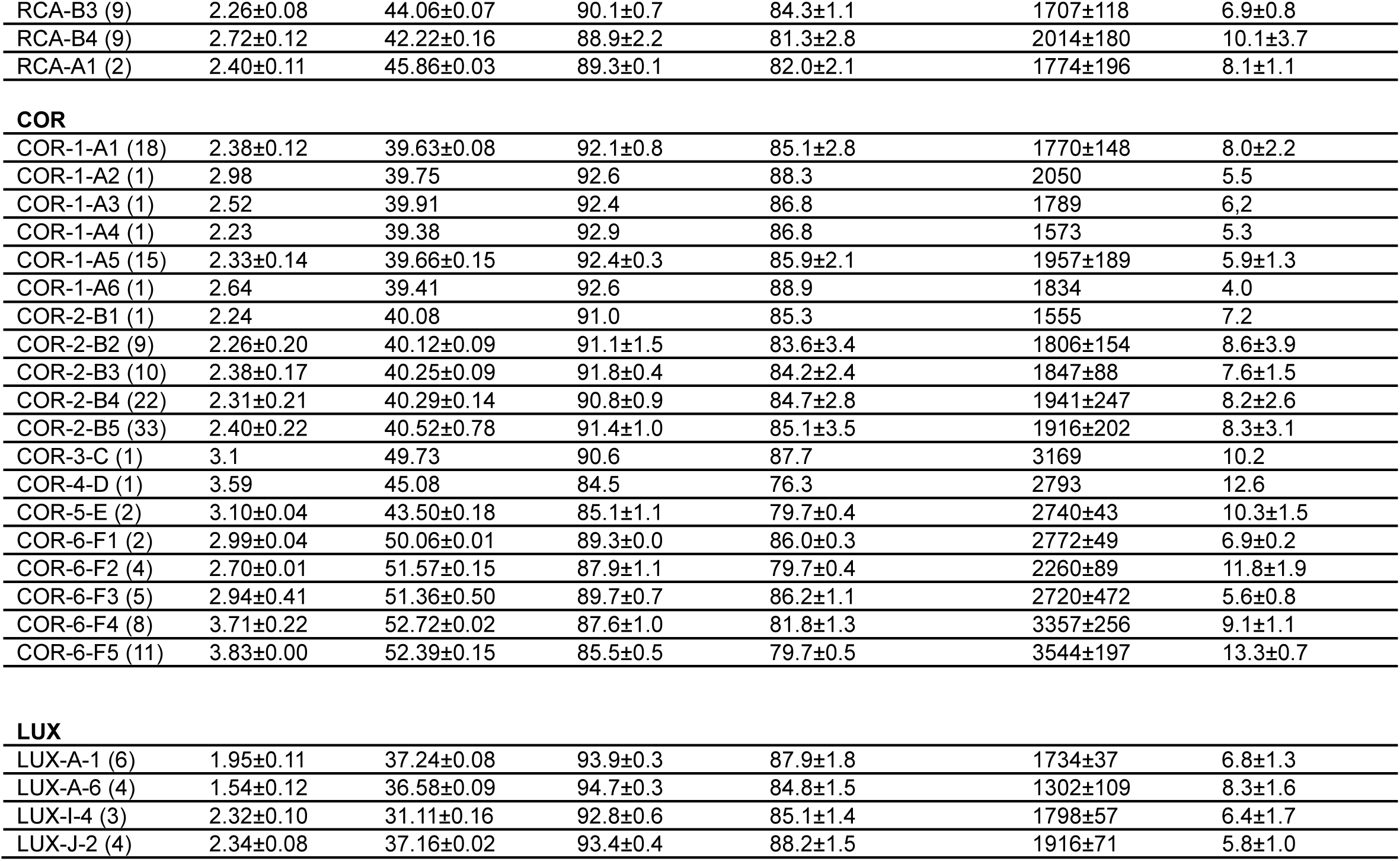
Genomic characteristics of species of the AAPR, AAR, TCR, RCA, COR, and LUX clusters of the *Roseobacter* group, including species identification, number of genomes per species (MAG/SAGs/isolates, in parenthesis), genome size ± standard deviation (SD), G+C content ± SD, coding density ± SD, coding density of intact genes± SD, and number of coding sequences (CDS number) ± SD. For the LUX cluster data are only presented for species represented by at least 3 MAGs or SAGs.

| Species | Genome Size (Mbp) | G+C Content (%) | Coding Density (%) | Coding Density (%) of intact genes ± SD | CDS number | Pseudogenes (% of CDS) |
| --- | --- | --- | --- | --- | --- | --- |
| **AAPR** |  |  |  |  |  |  |
| AAPR-A (13) | 2.41±0.10 | 37.59±0.04 | 92.9±0.4 | 85.1±2.4 | 1988±94 | 7.7±2.0 |
| AAPR-B (9) | 2.35±0.13 | 37.46±0.08 | 91.7±1.2 | 85.3±2.1 | 1832±77 | 7.5±2.1 |
| AAPR-C (16) | 2.22±0.08 | 37.05±0.06 | 91.4±1.2 | 84.0±2.1 | 1788±100 | 8.9±2.2 |
| **AAR** |  |  |  |  |  |  |
| AAR-A (1) | 2.57 | 38 | 92.6 | 84.8 | 2249 | 6.1 |
| AAR-B (1) | 2.48 | 37.91 | 91.1 | 83.1 | 2100 | 8.5 |
| AAR-C (2) | 2.30±0.07 | 38.10±0.04 | 91.7±0.3 | 86.1±4.0 | 1872±168 | 5.1±2.5 |
| AAR-D (3) | 2.39±0.10 | 38.18±0.06 | 91.7±0.3 | 86.9±1.2 | 1986±112 | 5.9±3.1 |
| AAR-E (13) | 2.36±0.08 | 38.41±0.06 | 90.6±0.8 | 82.2±2.3 | 1850±110 | 8.4±2.1 |
| **TCR** |  |  |  |  |  |  |
| TCR-A (3) | 2.13±0.04 | 38.89±0.08 | 89.4±0.9 | 85.6±2.4 | 1686±217 | 8.0±2.3 |
| TCR-B (17) | 2.37±0.08 | 38.60±0.09 | 90.2±0.2 | 88.3±1.2 | 2114±69 | 3.9±0.9 |
| TCR-C (3) | 2.28±0.18 | 36.54±0.34 | 92.6±0.3 | 87.2±0.53 | 1828±149 | 5.5±0.6 |
| TCR-D (12) | 2.26±0.17 | 36.64±0.21 | 91.8±1.3 | 86.2±3.5 | 1784±211 | 6.9±4.0 |
| TCR-E (29) | 2.17±0.12 | 36.02±0.08 | 92.6±0.2 | 89.4±1.7 | 1882±87 | 3.6±1.3 |
| TCR-F (1) | 2.65 | 48.29 | 91.5 | 88.6 | 2713 | 5.2 |
| TCR-G (2) | 2.61±0.01 | 50.17±0.01 | 92.1±0.2 | 90.2±0.7 | 2520±11 | 3.5±0.2 |
| **RCA** |  |  |  |  |  |  |
| RCA-C1 (3) | 2.45±0.13 | 48.56±0.15 | 90.3±0.2 | 80.5±1.6 | 1754±170 | 9.2±0.7 |
| RCA-C2 (1) | 3.12 | 46.36 | 86.4 | 77.2 | 2122 | 13.0 |
| RCA-C3 (14) | 2.64±0.21 | 52.00±0.49 | 89.9±0.7 | 85.6±1.5 | 2209±295 | 10.0±1.6 |
| RCA-C4 (18) | 2.85±0.19 | 47.91±0.11 | 89.2±0.3 | 85.2±1.0 | 2144±131 | 6.8±0.7 |
| RCA-C5 (7) | 3.12±0.18 | 47.97±0.122 | 89.1±0.4 | 83.7±2.7 | 2426±90 | 8.2±1.9 |
| RCA-C6 (6) | 3.09±0.29 | 53.70±0.93 | 90.2±1.3 | 85.6±2.0 | 3040±330 | 10.3±2.9 |
| RCA-C7 (2) | 2.49±0.01 | 53.56±0.03 | 90.0±0.7 | 81.8±0.9 | 2104±60 | 15.3±1.7 |
| RCA-C8 (1) | 2.92 | 53.22 | 89.8 | 83.9 | 2112 | 11.9 |
| RCA-B1 (2) | 2.36±0.17 | 43.69±0.06 | 89.7±0.0 | 82.7±2.2 | 1844±37 | 7.9±1.1 |
| RCA-B2 (23) | 2.34±0.11 | 43.72±0.11 | 90.8±0.3 | 87.3±1.7 | 2042±108 | 4.9±1.0 |
| RCA-B3 (9) | 2.26±0.08 | 44.06±0.07 | 90.1±0.7 | 84.3±1.1 | 1707±118 | 6.9±0.8 |
| RCA-B4 (9) | 2.72±0.12 | 42.22±0.16 | 88.9±2.2 | 81.3±2.8 | 2014±180 | 10.1±3.7 |
| RCA-A1 (2) | 2.40±0.11 | 45.86±0.03 | 89.3±0.1 | 82.0±2.1 | 1774±196 | 8.1±1.1 |
| **COR** |  |  |  |  |  |  |
| COR-1-A1 (18) | 2.38±0.12 | 39.63±0.08 | 92.1±0.8 | 85.1±2.8 | 1770±148 | 8.0±2.2 |
| COR-1-A2 (1) | 2.98 | 39.75 | 92.6 | 88.3 | 2050 | 5.5 |
| COR-1-A3 (1) | 2.52 | 39.91 | 92.4 | 86.8 | 1789 | 6.2 |
| COR-1-A4 (1) | 2.23 | 39.38 | 92.9 | 86.8 | 1573 | 5.3 |
| COR-1-A5 (15) | 2.33±0.14 | 39.66±0.15 | 92.4±0.3 | 85.9±2.1 | 1957±189 | 5.9±1.3 |
| COR-1-A6 (1) | 2.64 | 39.41 | 92.6 | 88.9 | 1834 | 4.0 |
| COR-2-B1 (1) | 2.24 | 40.08 | 91.0 | 85.3 | 1555 | 7.2 |
| COR-2-B2 (9) | 2.26±0.20 | 40.12±0.09 | 91.1±1.5 | 83.6±3.4 | 1806±154 | 8.6±3.9 |
| COR-2-B3 (10) | 2.38±0.17 | 40.25±0.09 | 91.8±0.4 | 84.2±2.4 | 1847±88 | 7.6±1.5 |
| COR-2-B4 (22) | 2.31±0.21 | 40.29±0.14 | 90.8±0.9 | 84.7±2.8 | 1941±247 | 8.2±2.6 |
| COR-2-B5 (33) | 2.40±0.22 | 40.52±0.78 | 91.4±1.0 | 85.1±3.5 | 1916±202 | 8.3±3.1 |
| COR-3-C (1) | 3.1 | 49.73 | 90.6 | 87.7 | 3169 | 10.2 |
| COR-4-D (1) | 3.59 | 45.08 | 84.5 | 76.3 | 2793 | 12.6 |
| COR-5-E (2) | 3.10±0.04 | 43.50±0.18 | 85.1±1.1 | 79.7±0.4 | 2740±43 | 10.3±1.5 |
| COR-6-F1 (2) | 2.99±0.04 | 50.06±0.01 | 89.3±0.0 | 86.0±0.3 | 2772±49 | 6.9±0.2 |
| COR-6-F2 (4) | 2.70±0.01 | 51.57±0.15 | 87.9±1.1 | 79.7±0.4 | 2260±89 | 11.8±1.9 |
| COR-6-F3 (5) | 2.94±0.41 | 51.36±0.50 | 89.7±0.7 | 86.2±1.1 | 2720±472 | 5.6±0.8 |
| COR-6-F4 (8) | 3.71±0.22 | 52.72±0.02 | 87.6±1.0 | 81.8±1.3 | 3357±256 | 9.1±1.1 |
| COR-6-F5 (11) | 3.83±0.00 | 52.39±0.15 | 85.5±0.5 | 79.7±0.5 | 3544±197 | 13.3±0.7 |
| **LUX** |  |  |  |  |  |  |
| LUX-A-1 (6) | 1.95±0.11 | 37.24±0.08 | 93.9±0.3 | 87.9±1.8 | 1734±37 | 6.8±1.3 |
| LUX-A-6 (4) | 1.54±0.12 | 36.58±0.09 | 94.7±0.3 | 84.8±1.5 | 1302±109 | 8.3±1.6 |
| LUX-I-4 (3) | 2.32±0.10 | 31.11±0.16 | 92.8±0.6 | 85.1±1.4 | 1798±57 | 6.4±1.7 |
| LUX-J-2 (4) | 2.34±0.08 | 37.16±0.02 | 93.4±0.4 | 88.2±1.5 | 1916±71 | 5.8±1.0 |

MAGs of the TCR cluster were recruited from ocean regions of temperate to polar latitudes (Supplementary Fig. S3A, Supplementary Table S4). MAGs of species TCR-A originate exclusively from the Southern Ocean and those of species TCR-B and -G from the Arctic Ocean.

MAGs of species TCR-D and -E were also recruited from the Arctic Ocean, but in addition from the temperate Atlantic Ocean, the North Sea and the genome of *Rhodobacterales* bacterium HTCC2255 from the eastern temperate Pacific. The single MAG of species TCR-F was recruited from the North Sea whereas the three MAGs of species TCR-C from the temperate Atlantic, the North Sea and the western Mediterranean Sea.

### AAR cluster

Twenty MAGs affiliate to the AAR cluster, classified into five distinct species represented by single and up to 13 MAGs (Fig. 2A, Table 1, Supplementary Fig. S3, Supplementary Tables S6 and S7). The AAR cluster exhibits relatively small genome sizes ranging from 2.24 to 2.57 Mbp (mean 2.38 ± 0.09 Mbp), low CDS counts ranging from 1681 to 2249 (mean 1905.15 ± 148.15), a low G+C content (mean 38.30 ± 0.17%) and a high coding density (mean of all gene features 90.96 ± 0.88%; mean for intact genes 83.44 ± 2.87%).

All MAGs of species AAR-E were recovered from the Arctic Ocean, whereas MAGs of the other four species from the temperate North and South Atlantic between 31° and 54° absolute latitude and the North Sea (Supplementary Fig. S3A, Supplementary Table S6).

### AAPR cluster

The AAPR cluster comprises 37 MAGs and one SAG, classified into three species (Fig. 2A, Table 1, Supplementary Fig. S3, Supplementary Tables S8 and S9). Their genomic features are not significantly different and rather similar to the AAR cluster with a mean genome size of 2.32 ± 0.13 Mbp, a mean number of CDS of 1866.82 ± 127.90, a mean G+C content of 37.33 ± 0.26% and a coding density of 91.98 ± 1.17% of all gene features and of intact genes of 84.57 ± 2.29 (Fig. 2B, Supplementary Table S8).

Recruitment locations exhibit distinct biogeographic patterns (Supplementary Fig. S3A, Supplementary Table S8). MAGs of species AAPR-A were recruited from the temperate North and South Atlantic between 31° and 54° absolute latitude. MAGs of species AAPR-B were also recruited from the temperate North and South Atlantic between 35° and 54.6° absolute latitude but in addition from the temperate North and South Pacific in the Californian and Peru upwelling regions. All MAGs of species AAPR-C originated from the Arctic Ocean except one which was recruited from the Patagonian Shelf of the South Atlantic.

### RCA cluster

The RCA cluster consists of three genera and 13 species represented by 83 MAGs and five isolates (Fig. 2A, Table 1, Supplementary Table S10). The eight species of the genus *Planktomarina* (RCA species C1-C8) exhibit genome sizes of 2.44 to 3.12 Mbp, CDS counts of 1754 to 3040 and G+C contents of 46.36% to 53.07% (Fig. 2B). In contrast, the four species of the genus *Pseudoplanktomarina* (RCA species B1-B4) and the MAGs of the genus *Cand*. Paraplanktomarina (RCA species A1, A2) have smaller genomes, 2.26 to 2.72 Mbp, CDS counts of 1708 to 2024 and G+C contents of 42.22% to 44.10%. Several genomic features among species of the *Planktomarina* and *Pseudoplanktomarina* genera were significantly different (Fig. 2B). MAGs were retrieved from the temperate Atlantic, Arctic and Southern Ocean and the North Sea. For further details see reference [7].

### COR cluster

The COR cluster is the largest of the six clusters, exhibiting the greatest diversity in genomic features (Fig. 2, Table 1, Supplementary Fig. S3, Supplementary Tables S11, S12). One hundred thirty MAGs, 14 SAGs and the genomes of two isolates (strains HIMB11 and SB2) were assigned to this cluster. It is subdivided into six subclusters which can be considered as genera based on < 70% ANI and comprises 19 species (based on ≥ 95% ANI, Fig. 2A, Supplementary Fig. S3).

The COR cluster shows a broad range of genomic characteristics: Genome sizes of individual species range from 2.23 to 3.83 Mbp, the G+C content from 39.38% to 52.72%, the coding density of all gene features from 85.06% to 92.94%, that of intact genes from 76.25% to 88.90%, and the number of CDS from 1555 to 3544 (Fig. 2B, Table 1, Supplementary Table S11).

Subclusters COR-1 and -2, encompassing six and five species, respectively, exhibit consistently features of genome streamlining regarding genome size, G+C content, coding densities of all and intact genes (Fig. 2B, Table 1, Supplementary Table S11). Subcluster COR-3 is represented only by the genome HIMB11, subcluster COR-4 by one MAG and subcluster COR-5 by one species with two MAGs. Subcluster COR-6 comprises five species and each species two to eleven MAGs. Species COR-6-E2 includes the genome of strain SB2 (Fig. 2A, Supplementary Table S11). Genomes of subclusters COR-3 to COR-6 are significantly larger, the CDS counts and G+C content significantly higher and coding densities of all genes significantly lower than those of subclusters COR-1 and -2 (Fig. 2B, Table 1, Supplementary Table S11).

The recruitment patterns of the different subclusters and genera of the COR cluster add interesting data to their genomic features. MAGs of subclusters COR-1 and -2 were recruited predominantly in the tropical and subtropical regions of the Atlantic, Pacific and Indian Ocean and only a few MAGs in temperate regions of the Mediterranean Sea, the Atlantic or Pacific Ocean (Supplementary Fig. S3A, Supplementary Table S11). Recruitment regions of the MAGs, genomes and species of the other subclusters were more variable and included the tropical central Pacific, the subtropical and temperate Atlantic Ocean, except MAGs of species COR-6-F2 which were exclusive to the Mediterranean Sea. MAGs of species COR-6-F5 were recruited from the North Sea, the temperate south Atlantic, the Arctic and the Southern Ocean (Supplementary Table S11). Two strains, not included in our analyses, affiliated to genome SB2 and a closely related species, and thus members of cluster COR-6-F3, were isolated in the subtropical eastern coastal Pacific Ocean at the San Pedro Ocean Time Series and in the East China Sea [58,59] thus expanding the recruitment regions of this species,

### LUX cluster

This cluster encompasses 27 SAGs and 19 MAGs classified into ten subclusters (Supplementary Tables S13 and S14). The subclusters encompass in total 30 species (Supplementary Fig. S3). However, 24 species are represented by only one or two MAGs or SAGs and only four species by at least four MAGs or SAGs, making an analysis of the within cluster diversity preliminary. The genome size of the entire cluster ranges between 1.45 and 2.82 Mbp, CDS counts between 1196 and 2323, the G+C content between 28.6% and 38.7% and the coding density between 90.5% and 95.2% for all genes and between 80.1% and 91.3% for intact genes (Supplementary Table S13). These genomic features of the four species comprising at least four MAGs or SAGs exhibited some significantly different characteristics such as genome size, G+C content, and CDS counts (Table 1). Our analysis is an extension of previous analyses which have already shown data on genomic features of this cluster (*Cand*. Luxecamonaceae family) [23,24].

The MAGs and SAGs of this cluster were recruited from tropical to temperate regions of the Atlantic, Pacific and Indian Oceans and the Red Sea, Mediterranean Sea and North Sea (Supplementary Table S13). All except two MAGs from the North Sea and Mediterranean Sea originated from locations within 40° absolute latitude.

### Genome streamlining in the *Roseobacter* clusters as compared to other pelagic prokaryotic groups

Our analyses show that each of the six pelagic clusters exhibits features of genome streamlining even though with distinct differences. Within the TCR cluster, the most deeply branching species F and G have larger genomes, higher CDS counts and a higher G+C content than the other five species of this cluster and the deeply branching AAR, AAPR and LUX clusters (Table 1, Fig. 2B). This indicates that these four clusters underwent genome streamlining relative to other deeply branching genera of the *Roseobacter* group and five TCR species even further within this cluster. The RCA cluster, branching off at an intermediate region of the phylogenetic tree of the *Roseobacter* group close to the genus *Nereida*, exhibits features of genome streamlining and expansion. The reduced genome size of species B1, B2, and B3 of the genus *Pseudoplanktomarina* and the lower G+C content of all four species of this genus relative to the most deeply branching RCA genus *Cand*. Paraplanktomarina indicate genome streamlining (Table 1). In contrast, the species of the genus *Planktomarina* appear to have undergone genome expansion and relaxation of growth under nutrient limitation resulting in larger genomes and a higher G+C content relative to *Cand*. Paraplanktomarina. For further details see reference [7]. Within the COR cluster, branching off further above the RCA cluster in the phylogenetic tree close to the genera *Pseudaestuariivita* and *Marinovum* (Fig. 1B, Supplementary Fig. S4), the species of subclusters COR-1 and -2 have smaller genomes, lower CDS counts and a lower G+C content than subclusters COR-3 to -6 (Table 1, Fig. 2B). Most species of the latter subclusters have also a lower coding density and a lower proportion of intact genes than subclusters COR-1 and COR-2. Hence, genome streamlining within the *Roseobacter* group happened not only at the deeply branching clusters but also in sublineages at intermediate regions of the phylogenetic tree, within the RCA and COR clusters. Another pelagic *Roseobacter* lineage with some genome streamlining features, CHUG, a genome size of 2.5 Mbp but a G+C content of 55%, has been recently identified [60]. Our results expand and refine the modelling study by Luo et al.[19] and other previous studies [7,60] and provide a solid basis to underscore that genome streamlining is an important evolutionary process within the *Roseobacter* group yielding prominent clusters and subclusters well adapted to energy- and nutrient-limited oceanic regions.

In contrast to the *Roseobacter* group the SAR11 clade/*Pelagibacterales*, the SAR86 clade and *Prochlorococcaceae* encompass genome sizes of < 3.1 Mbp and genome streamlining features of the entire order or major sublineages. The SAR11 and SAR86 clades exhibit genome sizes of < 1.4 and 1.7 Mbp and G+C contents of < 30% and < 36.5%, respectively, of the entire clade [20,26,61]. *Prochlorococcaceae* are subdivided into the low light adapted subfamily with genome sizes ranging between 1.7 and 2.7 Mbp and G+C contents of 34-51%, and the high light adapted subfamily with genome sizes of 1.6-1.8 Mbp and a G+C content of ∼31%. This cyanobacterial family with genome sizes smaller than of many other cyanobacterial families and restricted to the tropical and subtropical oceans, exhibits clear-cut adaptations to the high light and low nutrient regime in the near surface ocean by genome streamlining features. The SAR116 clade/*Puneispirillales* are subdivided into a high G+C (38-51%) and a low G+C (29-31%) subclade [21]. The genome sizes of the low G+C sublineages range from 1.7 to 2.7 Mbp and that of the high G+C sublineages from 2.1 to 3.1 Mbp. The SAR202 clade (*Chloroflexota*) is also structured into subgroups with streamlined genomes of 1.1-2 Mbp and G+C contents of 29-40% and other groups with genome sizes of 1.4-5 Mpb but G+C contents of 41-69% [32]. Thus, the latter two clades are also structured into subgroups with and without genome streamlined features, similar to the *Roseobacter* group.

A low number or proportion (< 11%) of pseudogenes and a high coding density (> 90%) have been discussed as indicative of genome streamlining in taxa with a high effective population size such as the SAR11 clade [20,61]. The TCR, AAR, AAPR and LUX clusters exhibit coding densities of all gene features of 89.4-94.7% and proportions of pseudogenes of 3.5-8.9% (Table 1). The RCA cluster features coding densities of 86.4-90.8% and proportions of pseudogenes of 4.9-15.3% and the COR cluster coding densities of 84.5-92.9% and proportions of pseudogenes of 4.0-13.3% (Table 1). Hence, the high coding densities and relatively low proportions of pseudogenes of the TCR, AAR, AAPR and LUX clusters suggest that these features were a result of genome streamlining, in line with the other features of genome streamlining of these clusters (see above). For the RCA cluster proportions of pseudogenes are more variable and generally higher. The higher coding densities of the COR-1 and COR-2 subclusters relative to the other COR subclusters are presumably a result of genome streamlining and purifying selection and are also in line with other features of genome streamlining of these subclusters (see above).

The number and proportion of pseudogenes as an indicator of genome streamlining, however, may need to be considered more cautiously and in the environmental context of the respective prokaryote. Copiotrophic and pathogenic bacteria encode also pseudogene proportions of only 2-7.5% [62,63], similar proportions as pelagic marine bacteria with streamlined genomes. Pseudogenes are suggested to arise from disablement of detectable native duplications, the decay of native single-copy genes, and failed horizontal transfers [63]. As pseudogenes may still act as regulators for gene transcription [62], the impact of purifying selection may vary accordingly such that the proportion of pseudogenes may not only be affected by genome streamlining.

### Functional features of the pelagic *Roseobacter* clusters

#### Energy metabolism

Proteorhodopsin (PR) is encoded in all species of the AAR and AAPR clusters, in species A to E of the TCR cluster, in species COR-5-E and in three of the five species of subcluster COR-6F (Figs. 1B, 3A, 4A, Supplementary Table S15). Nine of 13 RCA species encode PR as well (Supplementary Table S15) [7]. As the two most deeply branching species F and G of the TCR cluster and that of the RCA cluster, *Cand*. Paraplanktonmarina, do not encode the PR gene this mode of complementary light energy acquisition must have been gained via horizontal gene transfer. The same is true for the COR subclusters. To acquire the PR gene via horizontal gene transfer is rather common among bacterial lineages [64]. A phylogenetic analysis shows that the PR genes of the TCR, RCA and the COR-6-F affiliated species are rather closely related (Fig. 3A). The PR gene of the AAR and AAPR cluster and of species COR-5-E are located on different branches of this tree whereas that of the AAPR cluster is most closely related to those of the SAR11 and SAR86 clades. Our phylogenetic analysis revealed that PR is also encoded in a few species of other genera of the *Roseobacter* group not known before to carry this trait including *Ascidiaceihabitans*, *Yoonia* and *Roseovarius* (Figs. 1B and 3A, Supplementary Table S15). Another rhodopsin, xanthorhodopsin, has been detected in the genus *Octadecabacter* of the *Roseobacter* group [65] and we found it encoded also in MAGs of the genera *Ascidiaceihabitans* and *Yoonia* but not in the pelagic *Roseobacter* clusters (Supplementary Table S15, Supplementary Fig. S5A). All PR genes of the RCA and TCR clusters and the great majority of those of the AAR, AAPR and the COR clusters encode PR absorbing green light (Fig. 3A). Only few species of the latter three clusters encode PR absorbing blue light.

**Figure 3.**
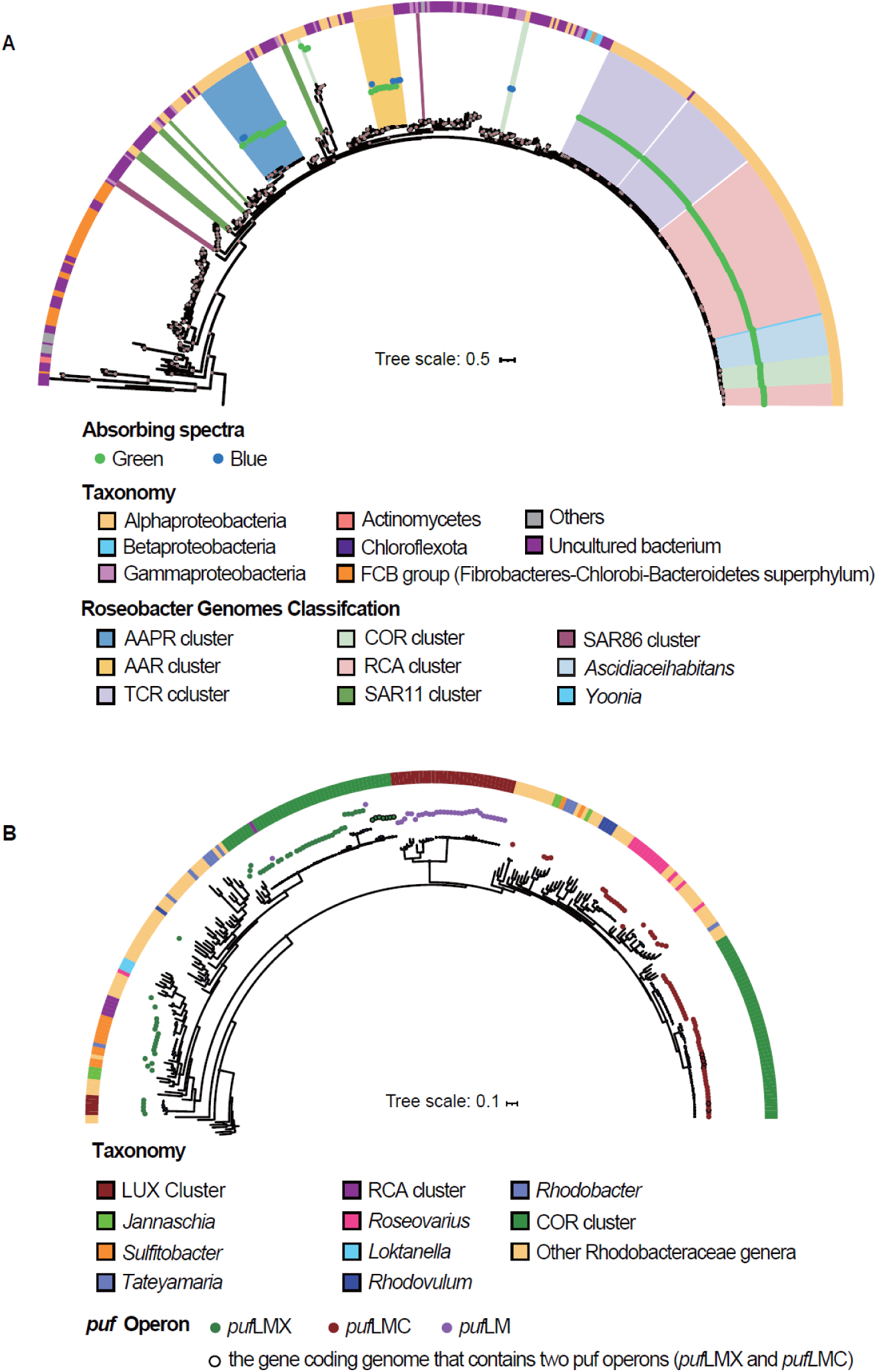
Phylogenetic trees of *puf*M and PR genes in the *Roseobacter* group. (A) The phylogenetic tree of the PR gene is based on 207 amino acid sequences from *Roseobacter* genomes of this study and 514 reference sequences from the UniProt database. Light absorption properties are indicated in blue and green dots. Outer semicircle colors represent genome classifications. Clusters and genera from this study and the SAR11 and SAR86 clades are highlighted by colored backgrounds. (B) The *puf*M phylogenetic tree is constructed from 100 *puf*M amino acid sequences from this study and 348 reference sequences. Color dots represent the operon types *puf*LMX, *puf*LMC and *puf*LM, identified in *Roseobacter* genomes. Seven COR cluster genomes harbor both *puf*LMX and *puf*LMC operons. Outer semicircle colors represent genome classifications.

**Figure 4.**
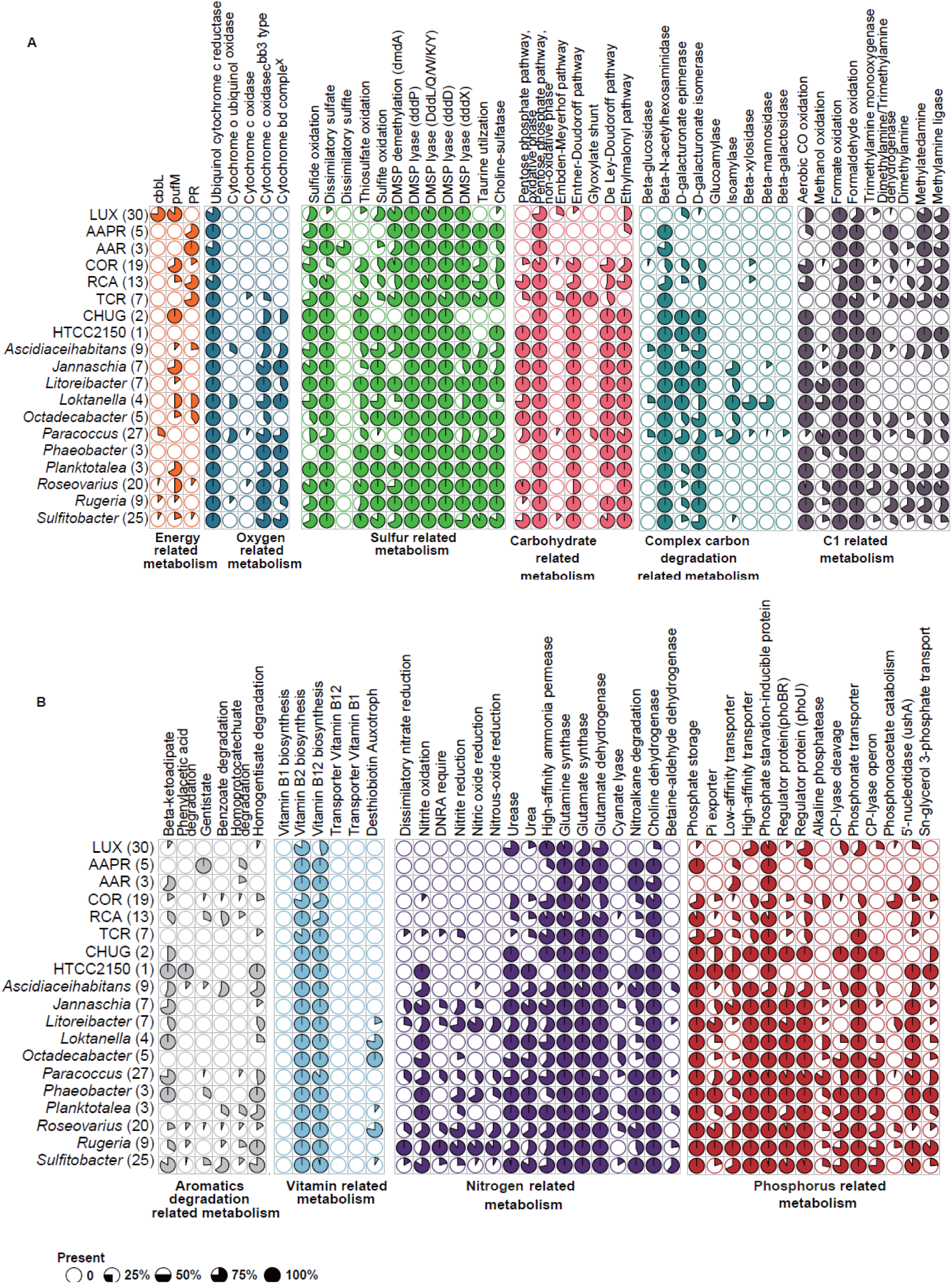
Functional features of pelagic *Roseobacter* cluster and representative genera. A metabolic module, pathway, or transporter is considered present in a genome if it reaches at least 75% completeness, as estimated using KEGG-Decoder and METABOLICS. A species is considered to possess a given feature if it is present in more than half of its genomes analyzed. The number of species within each cluster or genus included in this study is indicated in parentheses. Pie charts represent the proportion of species within each cluster or genus that contains the specified module, pathway or transporter. (A) Functional features associated with the metabolism of carbohydrates, complex carbon degradation, C1-compounds, nitrogen, and phosphorus. (B) Functional features associated with energy metabolism, oxygen utilization, vitamin biosynthesis, degradation of aromatic compounds, and sulfur metabolism. For detailed species-level information within the pelagic clusters, see Supplementary Tables S15–S24.

Complementary energy acquisition via PR is a common trait in the SAR11 and SAR116 clades, in sublineages of the SAR86 and SAR202 clades and several lineages of *Flavobacteriia* of *Bacteroidota* (Fig. 3A) [21,31,32,61,64]. PR can now be considered as of similar importance in pelagic *Roseobacter* clusters but exhibits a greater diversity than in other lineages as a result of several events of horizontal gene transfer.

The *Roseobacter* group is well known for AAP as an important trait of complementary light energy acquisition [16,66]. A phylogenetic analysis of the operons *puf*LM, *puf*LMC and *puf*LMX encoding the photosynthetic reaction center cytochrome c subunit revealed that it is not present in the TCR, AAPR, and AAR clusters but in species C6 to C8 of the genus *Plankto*-*marina* in the RCA cluster [7] and in 16 of the 19 species of the COR cluster including all but one species (represented only by one MAG) of the genome streamlined subclusters COR-1 and COR-2 (Figs. 1A, 3A, 4A, Supplementary Table S15). These operons have also been identified in the genomes of the LUX cluster [23,24] (Fig. 3B). Hence, AAP is important in but restricted to three of the six pelagic *Roseobacter* clusters, including the two rather diverse and genome streamlined subclusters COR-1 and COR-2 and the deeply-branching LUX cluster which is also able to fix CO_2_ [23]. The operon *puf*LMX exhibits a rather wide diversity among these clusters spanning almost the same width of the entire *Roseobacter* group (Fig. 3B). AAP has not been detected in other major pelagic bacterial lineages and thus is exclusive to the *Roseobacter* group. It has been shown for *Planktomarina temperata*, a member of the RCA cluster, that AAP is crucial for maintaining a higher cell abundance under nutrient starvation [67] and that this trait is transcribed during phytoplankton blooms [4,16].

Regarding other genetic features related to energy metabolism, the six pelagic *Roseobacter* clusters exhibit only minor differences compared to other genera of this group. Noteworthy is the absence of the genes encoding the cytochrome c oxidase cbb3 type and the cytochrome bd complex in all six clusters except the former in the two most deeply branching species F and G of the TCR cluster (Fig. 4A, Supplementary Table S16). Both genes encode cytochromes with a high affinity to oxygen and are used under reduced oxygen concentrations [68,69], traits obviously not required in pelagic systems with high oxygen concentrations.

Whereas the gene for sulfide oxidation (*sqr*) is encoded in most species of the six pelagic clusters thiosulfate oxidation is lacking in the AAR and AAPR clusters and in most species of the LUX cluster (Fig. 4A, Supplementary Table S17). Both oxidation pathways are common in most genera of the *Roseobacter* group [16,66]. The *sat* gene encoding dissimilatory sulfate reduction to sulfite is present in almost all species of the six pelagic clusters, except in subcluster COR-2, three species of subcluster COR-6 and in most species of the LUX cluster (Fig. 4A). However, the complementary gene encoding dissimilatory sulfate reduction to sulfite, *aprAB*, is exclusively encoded in the AAR cluster (Supplementary Table S17). It has been argued that the presence of both genes is essential for reducing sulfate to sulfite [70]. Hence, the potential to reduce sulfate to sulfite remains unclear in four of the five pelagic clusters.

#### Carbon, nitrogen, organic sulfur and phosphorous metabolism

It has been shown previously that the LUX cluster encodes the genes for CO_2_ fixation via the CBB cycle [23,24] (Fig. 4A). Regarding carbohydrate metabolism, the AAR, AAPR, and the LUX clusters are most limited among the six clusters. They lack the genes of the oxidative phase of the pentose phosphate pathway, the Entner-Doudoroff (ED) and the De Ley-Doudoroff pathways to metabolize glucose and galactose, respectively (Fig. 4A, Supplementary Table S18). This may explain why the AAR and AAPR clusters and the majority of species of the LUX cluster also lack the D-galacturonate isomerase and epimerase (Fig. 4A). All these pathways are encoded in almost all other genera of the *Roseobacter* group and the ED pathway has been shown to be the key pathway in marine bacteria to metabolize glucose [71]. Interestingly, the TCR cluster encodes the glyoxylate pathway but none of the other *Roseobacter* genera except a few species of *Paracoccus* (Fig. 4A, Supplementary Table S18). It has been shown that under severe substrate limitation a key gene of this pathway, isocitrate lyase, is highly expressed at light in the *Flavobacterium Dokdonia* spec. MED134 encoding PR and in the SAR11 clade [72]. Hence it is conceivable that the TCR cluster takes advantage of this pathway by saving carbon for biosynthetic requirements when light is available. On the other hand, the TCR cluster, as well as the AAR cluster, do not encode the ethylmalonyl pathway to convert acetyl-CoA to other metabolites for biosynthetic pathways, an alternative to the glyoxylate pathway [73] and present in all other genera of the *Roseobacter* group (Fig. 4A, Supplementary Table S18) [73].

The six clusters are quite versatile in using methylated amines as a nitrogen and energy source. Most species of these clusters are predicted to be able to metabolize monomethylamine, the AAPR cluster and several species of the AAR, TCR and COR clusters also trimethylamine and dimethylamine (Fig. 4A, Supplementary Table S19). The significance of these traits has been studied for model organisms of the *Roseobacter* group and documented also for this group in metagenomic studies [74,75], but has not been shown to be so widespread in pelagic *Roseobacter* lineages as in our study.

Decomposition of aromatic compounds is not common in the *Roseobacter* group [16,66]. Therefore, it is interesting that the gentisate pathway to break down hydroxybenzoate is encoded in all species of the AAPR cluster, in the four species of the genus *Pseudoplanktomarina* of the RCA cluster and in the COR-3 subcluster (Fig. 4B, Supplementary Table S20). It may enable these taxa to enter a niche not or little occupied by other pelagic bacterial groups in the epi- and mesopelagic to exploit an aromatic compound [76]. The *Roseobacter* group is deficient in the hydrolysis of polysaccharides, reflected by the absence of hydrolases of beta-linked polysaccharides in the six clusters and the great majority of other genera of the *Roseobacter* group (Fig. 4A). It is interesting, though, that the beta-N-acetyl hexosaminidase is encoded in most clusters and subclusters except the LUX cluster, a few species of the RCA cluster and the COR-1 and -2 subclusters (Fig. 4A, Supplementary Table S21), enabling them to utilize chito-oligosaccharides after breakdown to monomeric N-acetyl glucosamine.

The six pelagic clusters are predicted to have no or the TCR cluster very limited capabilities to reduce or oxidize inorganic nitrogen compounds, in line with most other genera of the *Roseobacter* group (Fig. 4B, Supplementary Table S22). Further, the AAPR and AAR clusters do not encode a urea transporter and urease but the high affinity ammonium permease even though it was not detected in all MAGs of these clusters (Fig. 4B, Supplementary Table S22).

Regarding the metabolism of organic sulfur compounds, the six pelagic clusters encode the features typical for the *Roseobacter* group, including the demethylation and cleavage pathways to degrade dimethylsulfonium propionate and taurine metabolism (Fig. 4A, Supplementary Table S17) [16,66]. The LUX cluster is unable to metabolize taurine. The pathways for the synthesis of vitamins B_2_, B_7,_ and B_12_ are present in the pelagic clusters, in line with most other genera of the *Roseobacter* group (Fig. 4B, Supplementary Table S23).

Phosphorous acquisition is variable among the six clusters and less versatile than in most other genera of the *Roseobacter* group (Fig. 4B, Supplementary Table S24). The TCR, AAR, AAPR and RCA clusters encode predominantly transporters for the high and/or low affinity phosphate transporter and the AAR and RCA clusters in addition the 5’-nucleotidase. Phosphonate transport is encoded in most species of four clusters but not in the AAPR and AAR clusters. It has been shown, though, that the phosphonate transporter PhnCDE can also transport phosphate [77] making it likely that this is also true for the pelagic *Roseobacter* clusters. The C-P lyase operon is absent except in 25% of the species of the LUX cluster. Subclusters COR-1 and -2 lack the high and low affinity phosphate transporter to a great extent and subclusters COR-3 to -6 the low affinity phosphate transporter (Supplementary Table S24). However, most species of these subclusters encode alkaline phosphatase, the phosphonoacetate hydrolase, the phosphonate transporter and quite a few also the C-P lyase. Hence the COR cluster is able to exploit organic phosphorous compounds, an important resource when inorganic phosphate is severely limiting or unavailable [78,79]. The LUX cluster is also predicted to be quite versatile in exploiting organic phosphorous compounds as most species encode several genes catalyzing the cleavage of organic phosphorous bonds (Fig. 4B, Supplementary Table S24). For further details on phosphorous acquisition by the clusters and within cluster differences see Supplementary Text.

#### Transporter proteins

Transporter proteins are very important for prokaryotes to successfully dwell in nutrient-limited pelagic environments, in particular SBP (solute binding protein) transporters comprising ABC, TRAP, TTT and TonB transporters. They include rather unspecific as well as highly specific transport proteins for compounds of different sizes and with different but also very high substrate affinities in the subnanomolar range [80,81]. Some of the SBP transporters have an even greater binding spectrum of substrates than deduced from earlier studies with model organisms [81]. Genes encoding SBP are among the most highly expressed genes of pelagic *Rhodobacterales* including MAGs closely related to members of the RCA (*Planktomarina*) and COR cluster (HIMB11) [82].

Because of their significance, we examined these features in the six *Roseobacter* clusters. SBP transporters comprised between 3.2% and 9.2% of total CDS (Table 2, Supplementary Fig. S6, Supplementary Tables S25-S30). In the AAR, AAPR and RCA cluster, percentages ranged from 5.5-7.3% with only little, if any statistically significant within cluster differences. Within the TCR cluster, however, SBP transporters of species TCR-A to -E ranged from 7.9-8.5% of total CDS whereas in species TCR-F and -G these transporters comprised only 4.1% and 5.1%, respectively, significantly lower proportions (Table 2, Supplementary Table S27). Similarly, the number of SBP transporters per Mbp was 41.7-49.7 in the latter two species whereas in the former five species 75.3-80.8 and thus significantly higher (Table 2, Supplementary Table S27). These differences indicate that SBP transporters in the TCR cluster were subjected less to genome streamlining or were even enriched in the streamlined species, possibly as an adaptation to the nutrient-limited environment in which they thrive. Interestingly, the COR cluster exhibited contrasting features. The most deeply branching species COR-6-F4 and -F5 exhibited the highest proportions of SBP transporters of total CDS, 8.4 ± 0.3% and 7.5 ± 0.2%, and also the highest numbers of SBP transporters per Mbp, 86.3 ± 2.7 and 78.9 ± 3.2, whereas the genome streamlined subclusters COR-1 and -2 exhibited proportions of 4.5-6.6% and numbers per Mbp of 43.4 to 66.1, values significantly lower than those of species COR-6-F1 and -F2 (Table 2, Supplementary Fig. S6, Supplementary Table S29). Obviously, these COR-subclusters adapted differently to the nutrient-limited environment than the TCR-species, by a more selective conservation of distinct SBP transporters.

**Table 2.** Transporter systems of species of the AAPR, AAR, TCR, RCA, COR, and LUX clusters of the *Roseobacter* group, including species identification, TonB, SBP, ABC, and TRAP transporters. Given are absolute numbers of transporters of each system ± standard deviation (SD), SBP and ABC transporters per Mbp, ABC transporters as a percentage of SBP transporters, and SBP transporters as a percentage of total CDS. For the LUX cluster data are only presented for species represented by at least 3 MAGs or SAGs

| Species | TonB transporters | SBP-dependent transporters | ABC transporters | TRAP transporters | TTT | ABC transporters (per Mbp) | SBP-dependent transporters (per Mbp) | ABC transporters (% SBP) | SBP-dependent transporters (% CDS) |
| --- | --- | --- | --- | --- | --- | --- | --- | --- | --- |
| **AAPR** |  |  |  |  |  |  |  |  |  |
| AAPR-A (13) | 1.07±0.28 | 115.2±7.8 | 110.0±7.9 | 4.2±1.0 | 1.0±0.0 | 52.1±2.6 | 54.5±2.9 | 95.5±0.9 | 5.8±0.3 |
| AAPR-B (9) | 0.11±0.33 | 109.8±6.8 | 106.0±6.3 | 2.4±0.9 | 1.3±0.7 | 54.3±3.3 | 56.2±3.6 | 96.6±0.8 | 6.0±0.4 |
| AAPR-C (16) | 0.94±0.25 | 108.6±9.9 | 106.3±9.2 | 1.8±1.2 | 0.5±0.5 | 55.2±3.4 | 56.4±3.6 | 97.0±1.3 | 6.1±0.4 |
| **AAR** |  |  |  |  |  |  |  |  |  |
| AAR-A (1) | 1 | 145 | 137 | 6 | 2 | 56.1 | 59.4 | 94.5 | 6.5 |
| AAR-B (1) | 2 | 120 | 113 | 6 | 1 | 50.3 | 53.4 | 94.2 | 5.7 |
| AAR-C (2) | 1.00±0.00 | 125.0±11.3 | 119.5±10.6 | 4.50±0.7 | 1.0±0.0 | 59.8±0.3 | 62.6±0.2 | 95.6±0.2 | 6.7±0.0 |
| AAR-D (3) | 1.00±0.00 | 129.7±14.3 | 120.7±12.1 | 4.33±1.5 | 4.7±1.2 | 56.8±3.1 | 61.0±3.6 | 93.1±1.4 | 6.5±0.4 |
| AAR-E (13) | 0.77±0.44 | 123.6±8.8 | 116.5±7.7 | 5.61±1.7 | 1.5±1.2 | 58.1±3.3 | 61.6±3.6 | 94.3±1.1 | 6.7±0.4 |
| **TCR** |  |  |  |  |  |  |  |  |  |
| TCR-A (3) | 0.67±0.58 | 137.7±15.9 | 122.3±14.4 | 10.0±0.0 | 5.3±1.5 | 68.3±1.5 | 76.9±2.0 | 88.8±0.4 | 8.2±0.3 |
| TCR-B (17) | 1.00±0.00 | 170.5±10.9 | 145.2±8.4 | 19.6±3.7 | 5.8±0.7 | 64.7±2.5 | 76.0±3.0 | 85.2±1.7 | 8.1±0.3 |
| TCR-C (3) | 1.00±0.00 | 155.7±22.4 | 139.3±17.0 | 11.0±4.6 | 5.3±1.5 | 72.5±4.8 | 80.8±5.7 | 89.7±2.4 | 8.5±0.6 |
| TCR-D (12) | 0.83±0.39 | 141.4±24.4 | 125.9±20.1 | 11.3±4.7 | 4.2±1.2 | 67.2±5.3 | 75.3±6.0 | 89.2±2.1 | 7.9±0.6 |
| TCR-E (29) | 1.00±0.00 | 148.9±15.6 | 135.8±13.9 | 11.0±2.5 | 3.1±0.9 | 68.3±4.6 | 75.5±5.1 | 90.6±1.4 | 7.9±0.5 |
| TCR-F (1) | 1 | 111 | 99 | 6 | 6 | 37.2 | 41.7 | 89.2 | 4.1 |
| TCR-G (2) | 2.00±0.00 | 128.0±1.4 | 113.0±1.4 | 13.0±0.0 | 2.0±0.0 | 43.8±0.8 | 49.7±0.9 | 88.3±0.1 | 5.1±0.1 |
| **RCA** |  |  |  |  |  |  |  |  |  |
| RCA-C1 (3) | 0.67±0.58 | 96.0±4.4 | 88.3±3.1 | 7.7±2.5 | 0.0±0.0 | 46.4±2.5 | 46.4±2.5 | 92.1±2.4 | 5.55±0.4 |
| RCA-C2 (1) | 1 | 122 | 111 | 11 | 0 | 47.8 | 52.5 | 91.0 | 5.8 |
| RCA-C3 (14) | 1.21±0.42 | 124.4±29.3 | 115.1±26.4 | 5.6±2.7 | 3.6±1.7 | 49.4±5.9 | 53.3±6.4 | 92.7±1.9 | 5.6±0.7 |
| RCA-C4 (18) | 1.00±0.00 | 131.3±8.6 | 121.3±8.3 | 10.0±0.5 | 0.0±0.0 | 52.8±3.6 | 57.1±3.7 | 92.4±0.4 | 6.1±0.4 |
| RCA-C5 (7) | 1.00±0.00 | 155.0±8.4 | 142.7±7.7 | 10.7±1.1 | 1.6±0.5 | 55.0±2.8 | 59.7±3.1 | 92.1±0.5 | 6.4±0.4 |
| RCA-C6 (6) | 1.00±0.00 | 185.0±21.9 | 163.7±17.4 | 15.3±3.2 | 6.7±2.7 | 57.9±4.1 | 54.9±3.4 | 88.6±2.2 | 6.1±0.2 |
| RCA-C7 (2) | 0.50±0.71 | 125.2±2.1 | 116.5±2.1 | 6.0±0.0 | 3.0±0.0 | 53.8±0.6 | 58.0±0.8 | 92.8±0.1 | 6.0±0.1 |
| RCA-C8 (1) | 1 | 121 | 107 | 11 | 3 | 49.1 | 55.5 | 88.4 | 5.7 |
| RCA-B1 (2) | 1.00±0.00 | 110.5±21.9 | 105.5±20.5 | 4.0±1.4 | 1.0±0.0 | 53.4±9.0 | 56.0±9.7 | 95.5±0.4 | 6.0±1.1 |
| RCA-B2 (23) | 1.00±0.00 | 118.1±13.4 | 111.6±12.3 | 5.3±0.8 | 1.3±0.9 | 51.2±3.3 | 54.2±3.5 | 94.5±0.9 | 5.8±0.4 |
| RCA-B3 (9) | 1.00±0.50 | 97.8±11.5 | 93.4±10.8 | 3.6±1.2 | 0.8±0.4 | 50.3±3.8 | 52.6±4.0 | 95.6±1.1 | 5.7±0.5 |
| RCA-B4 (9) | 0.67±0.71 | 146.6±22.3 | 139.9±21.1 | 5.8±1.9 | 0.9±0.3 | 63.3±4.8 | 66.3±5.2 | 95.5±0.9 | 7.3±0.6 |
| RCA-A1 (2) | 1.00±1.41 | 112.0±14.1 | 107.5±13.0.4 | 1.5±0.7 | 3.0±0.0 | 55.7±0.9 | 58.0±1.0 | 96.0±0.1 | 6.3±0.1 |
| **COR** |  |  |  |  |  |  |  |  |  |
| COR-1-A1 (18) | 1.83±0.83 | 109.9±11.4 | 104.8±10.7 | 2.3±1.6 | 2.8±0.7 | 56.6±4.6 | 59.3±4.7 | 95.4±1.5 | 6.2±0.5 |
| COR-1-A2 (1) | 1 | 115 | 112 | 2 | 1 | 53.5 | 54.9 | 97.4 | 5.6 |
| COR-1-A3 (1) | 1 | 117 | 115 | 1 | 1 | 62.1 | 63.2 | 98.3 | 6.5 |
| COR-1-A4 (1) | 2 | 71 | 70 | 0 | 1 | 42.7 | 43.4 | 98.6 | 4.5 |
| COR-1-A5 (15) | 0.93±0.26 | 114.9±17.5 | 112.7±16.9 | 1.3±0.9 | 0.9±0.5 | 55.4±3.9 | 56.4±4.1 | 98.2±0.7 | 5.8±0.4 |
| COR-1-A6 (1) | 1 | 108 | 104 | 3 | 1 | 54.7 | 56.9 | 96.3 | 5.9 |
| COR-2-B1 (1) | 0 | 91 | 88 | 3 | 0 | 54.9 | 56.8 | 96.7 | 5.9 |
| COR-2-B2 (9) | 1.00±0.00 | 115.8±13.7 | 109.7±13.5 | 3.9±1.7 | 2.2±0.7 | 57.6±3.9 | 60.8±4.0 | 94.7±0.8 | 6.4±0.4 |
| COR-2-B3 (10) | 1.00±0.00 | 124.5±5.9 | 118.0±5.5 | 4.6±1.4 | 1.9±0.7 | 61.1±2.4 | 64.4±2.6 | 94.8±1.1 | 6.8±0.3 |
| COR-2-B4 (22) | 1.09±0.29 | 135.1±25.8 | 127.9±23.9 | 5.0±2.0 | 2.5±0.8 | 62.1±5.7 | 65.7±6.4 | 94.5±1.1 | 6.9±0.6 |
| COR-2-B5 (33) | 0.94±0.24 | 132.6±17.3 | 125.9±15.6 | 4.6±1.7 | 2.0±0.9 | 62.9±4.1 | 66.1±4.5 | 95.1±1.2 | 6.9±0.4 |
| COR-3-C (1) | 1 | 162 | 147 | 7 | 8 | 47.4 | 52.3 | 90.7 | 5.1 |
| COR-4-D (1) | 1 | 200 | 191 | 5 | 4 | 62.1 | 65.0 | 95.5 | 7.2 |
| COR-5-E (2) | 1.00±0.00 | 148.0±8.5 | 142.0±5.7 | 4.0±1.4 | 2.0±1.4 | 48.6±0.7 | 50.6±1.6 | 96.0±1.7 | 5.4±0.2 |
| COR-6-F1 (2) | 1.00±0.00 | 190.0±1.4 | 177.0±4.2 | 9.5±2.1 | 3.5±0.7 | 60.8±2.2 | 65.3±1.3 | 93.2±1.5 | 6.9±0.2 |
| COR-6-F2 (4) | 1.00±0.00 | 146.7±8.1 | 138.0±7.8 | 4.3±1.3 | 4.5±0.6 | 56.7±2.6 | 60.2±2.3 | 94.0±1.1 | 6.5±0.3 |
| COR-6-F3 (5) | 1.00±0.00 | 210.6±44.8 | 195.4±39.8 | 9.8±3.8 | 5.4±1.5 | 67.8±4.0 | 73.0±4.8 | 92.9±0.9 | 7.7±0.5 |
| COR-6-F4 (8) | 1.00±0.00 | 309.8±27.3 | 280.8±23.6 | 19.4±3.3 | 9.3±2.3 | 78.2±2.4 | 86.3±2.7 | 90.7±1.0 | 9.2±0.3 |
| COR-6-F5 (11) | 1.00±0.00 | 293.4±24.1 | 266.5±19.7 | 19.3±1.8 | 7.6±3.3 | 71.7±2.5 | 78.9±3.2 | 90.9±0.8 | 8.3±0.3 |
| **LUX** |  |  |  |  |  |  |  |  |  |
| LUX-A-1 (6) | 1.00±0.00 | 71.3±7.5 | 68.8±7.4 | 2.5±0.5 | 0.0±0.0 | 39.7±4.3 | 41.1±4.3 | 96.5±0.9 | 4.1±0.4 |
| LUX-A-6 (4) | 2.00±0.00 | 42.5±8.3 | 40.5±8.3 | 0.0±0.0 | 2.0±0.0 | 30.7±4.0 | 32.3±3.9 | 95.1±1.2 | 3.2±0.4 |
| LUX-I-4 (3) | 2.00±0.00 | 128.0±13.0 | 118.0±12.4 | 6.0±0.0 | 4.0±1.0 | 63.0±3.3 | 68.3±3.4 | 92.2±0.6 | 7.1±0.5 |
| LUX-J-2 (4) | 1.00±0.00 | 110.5±13.0 | 107.8±5.4 | 1.8±0.8 | 1.0±0.0 | 5.6±0.2 | 55.5±2.4 | 97.48±1.0 | 5.8±0.2 |

ABC transporters constituted SBP transporters in all six clusters to at least 85%, in the AAPR, AAR and LUX clusters and in subclusters COR-1 and -2 even to >92% (Table 2). The lowest fractions of 85.2% to 90.6% were recorded in the TCR cluster. The number of TRAP transporters was generally much lower and did not exceed 19 in any cluster (Table 2, Supplementary Tables S25-S30). The number of TonB and TTT transporters was even lower and no species of any cluster encoded more than two and eight, respectively (Table 2, Supplementary Tables S25-S30). This shows the overwhelming significance of ABC transporters for the pelagic *Roseobacter* clusters, obviously reflecting their adaptation to the nutrient-poor pelagic environment. Previous analyses of transporters in marine epipelagic bacteria showed that in *Alphaproteobacteria* and more specifically in the SAR11 clade ABC transporters dominate whereas in *Bacteroidota* and *Gammaproteobacteria* TonB transporters [31,83,84].

To examine potential substrate profiles of the different pelagic clusters and the possible differences among species of a given cluster we identified categories of the SBP transporters according to the KEGG database. Among the saccharide, polyol and lipid transporter category only the rhamnose transporter is encoded in almost all species of five clusters but not in the LUX cluster of which only few species encode transporters for alpha-glycosides or sorbitol/mannitol (Fig. 5, Supplementary Table S31). In the AAR and AAPR clusters and the COR-1 and -2 subclusters the rhamnose transporter is the only encoded transporter of this category. The TCR, RCA and the other COR subclusters encode also several other transporters of this category such as for sorbitol/mannitol, multiple sugars, glucose/mannose and fructose. The pelagic clusters encode very limited potential capabilities for metallic cation and vitamin B_12_ transport and only few species of each of the six clusters encode manganese and iron transport and a few species of the COR cluster in addition transport of iron-siderophore (Fig. 5, Supplementary Table S32). Transporters of amino acids (branched-chain, general L-amino acids, polar amino acids) were encoded in most species of all clusters whereas transport of phosphate, phosphonate and urea was limited to a few species of the TCR, RCA COR, and LUX clusters (Fig. 4B, Fig. 5, Supplementary Table S33). Glycine betaine/proline transport was encoded in most species of five clusters but not in the AAR cluster (Fig. 5, Supplementary Table S34). The latter, in contrast to most species of the other clusters, encoded the iron (III) transporter. Transporters of the NitT/TauT family, specific for nitrate, citrate, cyanate, bicarbonate and taurine, aromatic sulfur, phthalate, hydroxymethylpyrimidine, were encoded in most species of the AAPR, AAR and COR clusters, two species of the LUX cluster but not in the TCR and RCA clusters (Fig. 4A, 5). Transporters for spermidine and putrescine were not encoded in the AAPR cluster but in most species of the other clusters (Fig. 5, Supplementary Table S34). Transporters for peptides/nickel and microcin C were also encoded in the great majority of species of all clusters but that for microcin C not in the AAPR, AAR clusters, the RCA *Cand*. Paraplanktomarina and *Pseudoplanktomarina* species and the COR subclusters COR-1 and -2 (Fig. 5, Supplementary Table S35).

**Figure 5.**
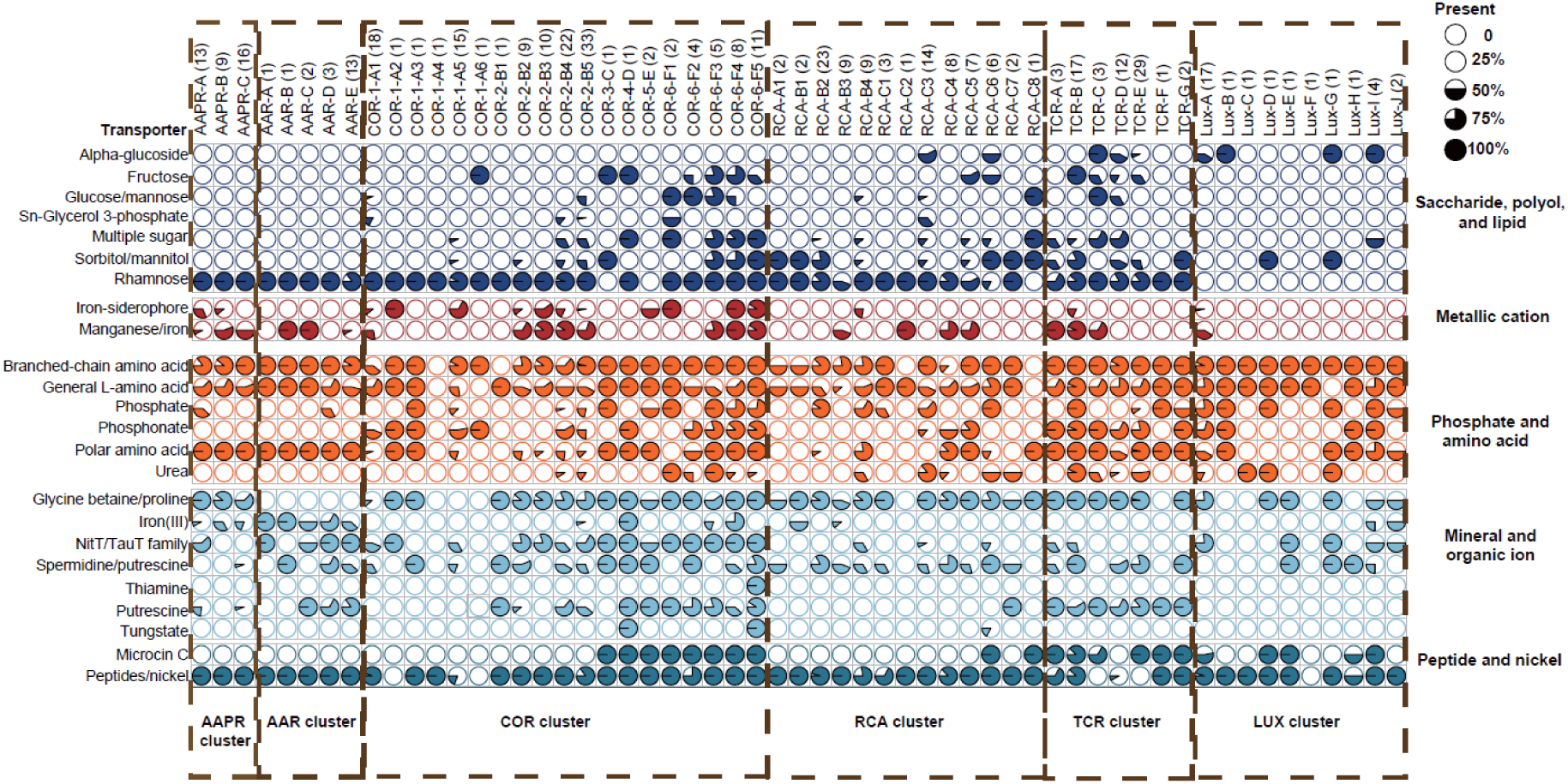
ABC transporter categories for species of the AAPR, AAR, TCR, RCA, COR, and LUX clusters. A transporter is considered present in a genome if at least 80% of its associated genes are detected. The number of genomes analyzed per species is indicated in parentheses; for the LUX cluster, the number of species per subcluster is indicated. Pie charts represent the proportion of genomes within each species that contains a specific transporter. For details of single species of the pelagic clusters see Supplementary Tables S25 to S35.

These transporter profiles indicate differences of the substrate preferences among the six pelagic *Roseobacter* clusters and their sublineages. However, amino acids, oligopeptides and polyamines were their major substrates. Transport and presumably subsequent metabolism of monosaccharides was more restricted, predominantly to species of the TCR, RCA and COR subclusters with larger genomes and thus less subjected to streamlining.

### Biogeography

To relate our data on genomic and functional features of the different clusters and species to their occurrence and abundance in different ocean regions and basins we mapped the reads of MAGs, SAGs and genomes of these clusters and species onto metagenomic reads of 745 samples encompassing the global oceans. Seventy percent of the samples originated from the epipelagic (< 200 m depth), 20% from the mesopelagic (200-1000 m) and 10% from the bathypelagic (> 1000 m). In the two latter layers the Mediterranean Sea and Red Sea were not represented.

The entire *Roseobacter* group constituted between < 2% and 24% of total reads in the epi- and mesopelagic and < 2-16% in the bathypelagic (Supplementary Fig. S7, Supplementary Tables S36 to S38). Highest percentages in the epipelagic of 15-22% were recorded in coastal areas, mainly in temperate regions, and in polar regions of both hemispheres. In the mesopelagic, highest percentages of 10-24% were recorded in all ocean regions and in the bathypelagic of 9-15.5% in all regions except the south tropical and north temperate region and the Southern Ocean (Supplementary Table S37 and S38). These data are in line with previous reports from different ocean regions and a focus on the epipelagic [5,85–89], but provide the first global overview on the relative abundance of this group of pelagic bacteria with a detailed extension to the meso- and bathypelagic.

In the epipelagic, the AAPR and TCR clusters were detected at relative abundances > 0.05% and up to 6.0% and 20.5% of total reads of the *Roseobacter* group, respectively, in the polar, temperate and subtropical regions of both hemispheres (Fig. 6A, Supplementary Table S39). The AAR cluster did not comprise more than 2.0% in any region. The RCA cluster constituted 17.2-44.6% in the temperate and polar regions of both hemispheres but < 3.2% in the other regions. The COR cluster constituted 18.8-57.2% from the north temperate to the south temperate regions but <4.4% in the polar regions of both hemispheres. The LUX cluster was even more restricted to temperate and warm waters and constituted 13.9-41.5% from the north temperate to the south subtropical regions, but only 3.8% in the south temperate and <0.3% in the polar regions of both hemispheres. Hence, the COR and LUX clusters dominated greatly the *Roseobacter* group from the north temperate to the south subtropical region, comprising 57-92%. All pelagic clusters together encompassed 85.6-93.2% from the north temperate to the south temperate regions and 53.6-67.2% of total reads of the *Roseobacter* group in both polar regions (Supplementary Table S39). Other genera, contributing substantial proportions in the latter two regions included *Ascidiaceihabitans*, EhC02, *Octadecabacter*, *Roseovarius*, *Sulfitobacter* and *Yoonia* (Fig. 6A, Supplementary Table S39).

**Figure 6.**
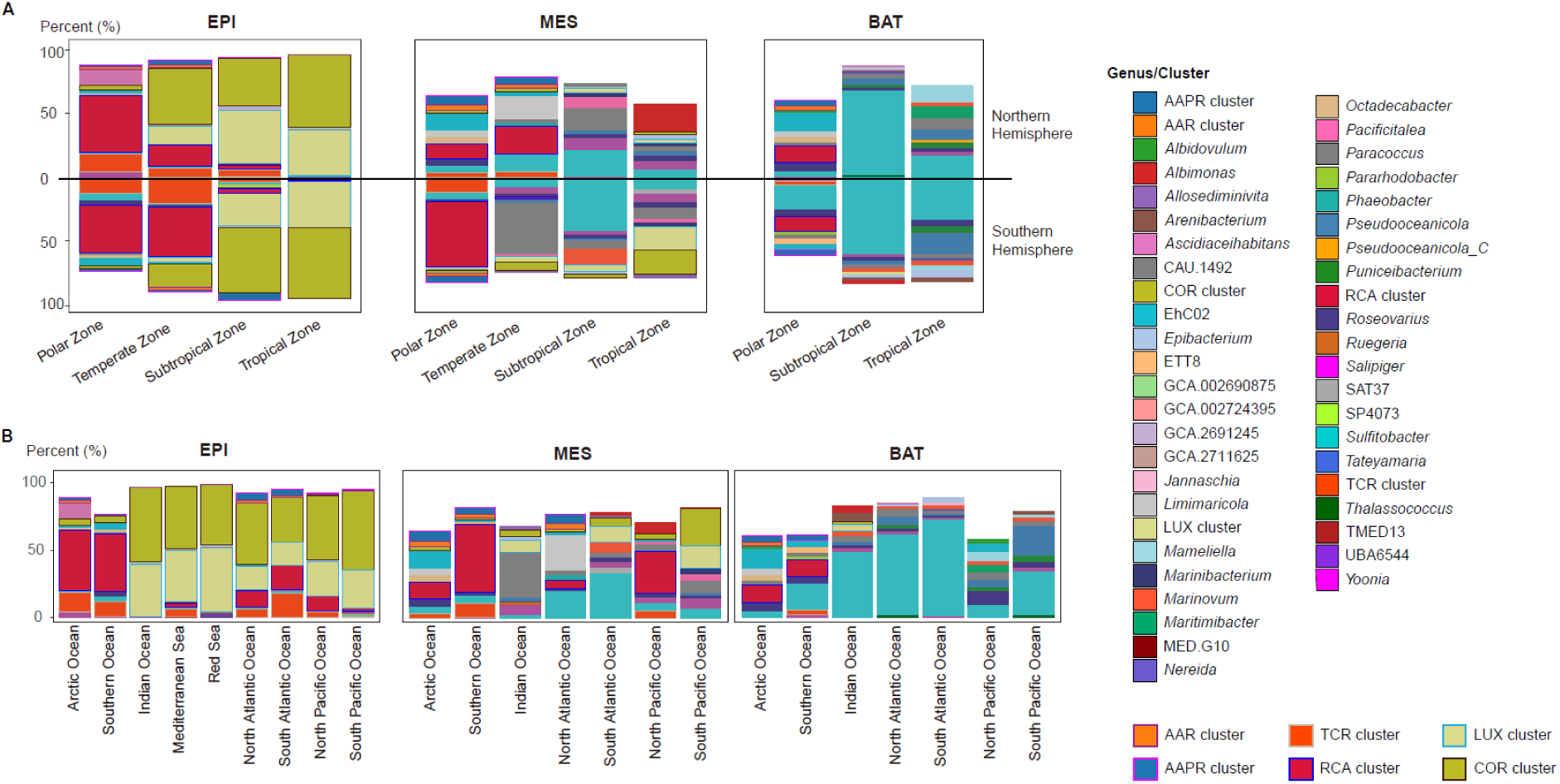
Biogeographic distribution of the pelagic clusters and genera of the *Roseobacter* group in the global oceans. (A) Relative abundances across different temperature zones: polar (latitude ≥ 66.5°), temperate (66.5° > latitude ≥35°), subtropical (35° > latitude ≥ 23.5°), tropical (latitude < 23.5°). In each panel, only the top 10 taxa by percentage are shown. Missing taxa would add up to 100%. (B) Relative abundances across different ocean regions in the epipelagic (EPI), mesopelagic (MES), and bathypelagic (BAT). For details on the different clusters and genera see Supplementary Tables S36 to S44.

The TCR, AAPR, AAR and RCA clusters were basically absent in the Indian Ocean and Red Sea (Fig. 6B, Supplementary Table S42) and the AAPR and AAR clusters constituted < 1.0% in the North and South Pacific. The TCR cluster constituted 10.5-18.2% in the Arctic, South Atlantic and Southern Ocean and the RCA cluster in the Arctic and Southern Ocean ∼43% and in the North and South Atlantic and the North Pacific 10.7-18.1% (Fig. 6A, Supplementary Table S42). The preference of the COR and LUX clusters to warmer waters was reflected by their high proportions in the warmer ocean basins. The COR cluster constituted 32.8-58.5% of all reads affiliated to the *Roseobacter* group in all ocean basins except the Arctic and Southern Ocean (Fig. 6B, Supplementary Table S42). The LUX cluster constituted 28.2-38.3% in the North and South Pacific, the Indian Ocean and the Mediterranean Sea and even 47.1% in the Red Sea. Our biogeographic patterns of the LUX cluster generally agree with those of previous studies (*Cand*. Luxescamonaceae family) but our more comprehensive dataset detected much higher percentages than previous studies [23,24].

There were also some distinct biogeographic epipelagic patterns of single species or subclusters. For example, species D of the AAR cluster was detected in all ocean regions whereas species E was only detected in the Arctic and Atlantic Ocean (Supplementary Fig. S8B). Species C of the AAPR cluster was not detected in the Mediterranean Sea and the South Pacific Ocean and species TCR-B was the only species of this cluster found in the Southern Ocean. All species of the genome streamlined subclusters COR-1 and -2 were detected only in the non-polar and predominantly in the tropical and subtropical regions whereas species of the other subclusters exhibited wider distribution patterns including the (sub)polar regions of both hemispheres. For further details of single species and subclusters see Supplementary Fig. S8 and Supplementary Text and for the biogeography of the RCA species reference [7].

In the mesopelagic, proportions of the pelagic clusters were substantially lower than in the epipelagic with percentages not exceeding 37.4% of total reads of the *Roseobacter* group except in the Southern Ocean where they constituted up to 70.1% (Fig. 6B, Supplementary Tables S40, S43). In the north subtropical and tropical and the south subtropical regions proportions did not exceed 4%. The AAPR cluster constituted < 0.4% from the north subtropical to the south subtropical region but 5.2-7.3% in the north temperate and both polar regions, proportions higher than in the epipelagic. Similarly, the AAR cluster constituted proportions of 2.3-4.4% in both polar and the north temperate regions, higher values than in the epipelagic. The TCR cluster exhibited proportions not exceeding 4.3% in all except the south polar region with a similar value as in the epipelagic of 10.4%. The RCA cluster was the most abundant pelagic cluster in the north temperate and both polar regions with the highest proportion of any pelagic cluster in the mesopelagic in the south polar region of 50.7% (Fig. 6). The COR cluster was of much lower significance in the meso-than in the epipelagic with proportions below 3.3% in all regions except the south tropical and south temperate region where it comprised 9.3% and 6.3%, respectively. The same was true for the LUX cluster with proportions of < 5.3% in all except the south tropical region where it constituted 18.3%. Other genera constituted relatively high proportions to the *Roseobacter* group community in the mesopelagic, including *Albimonas*, EhC02, *Marinovum*, *Paracoccus*, *Salipiger* and *Sulfitobacter*, which contributed up to 13-22% in different regions (Fig. 6, Supplementary Tables S40, S43). *Sulfitobacter* and *Marinovum* have been reported previously as prominent taxa of the prokaryotic community in the meso- and bathypelagic South Pacific and Atlantic Ocean [89,90].

In the bathypelagic, the pelagic clusters were of even lower significance than in the mesopelagic. Only in the polar regions of both hemispheres they accounted for about 22% of total reads of the *Roseobacter* group whereas in the other regions they did not exceed 1.6% (Fig. 6, Supplementary Tables S41 and S44). In the polar regions, the RCA and AAPR clusters constituted about 12% and 4% and the COR and LUX clusters < 1%. Other genera dominated the *Roseobacter* group. The genus *Sulfitobacter* was most prominent and comprised 33.7-67% in the north subtropical and south tropical to temperate regions and in other regions 4.5-17.6% (Fig. 5, Supplementary Tables S41 and S44). Other genera, including EhC02, *Mamiella*, *Maritimibacter*, *Pseudooceanicola* and *Roseovarius* were also prominent and constituted 10-22% in single regions (Fig. 6, Supplementary Tables S41 and S44). These findings indicate that the *Roseobacter* group and its pelagic clusters are prominent in the epi- and mesopelagic and adapted to the more productive and partially nutrient-limited upper layers of the global oceans whereas in the deep sea other prokaryotic lineages prevail.

### Biogeographic patterns related to genomic and functional features and conclusion

Our comprehensive analyses show interesting relationships between the biogeography in the epipelagic and genomic and functional features of the different clusters and cluster species. The deeply branching TCR, AAPR and AAR clusters, exhibiting small genomes and all except species TCR-G also a low G+C content, were detected only or predominantly in temperate and polar oceans. The same is true for the RCA cluster [7]. One species of the latter cluster and two species of the TCR cluster also occurred in the Mediterranean Sea and species AAR-D also in the Indian Ocean and Red Sea (Fig. 6, Supplementary Fig. S8, Supplementary Tables S4, S6, S8, S10). Most species of these clusters encode the PR gene, except the most deeply branching species TCR-G and species RCA-C6 to -C8 (Supplementary Table S15). The latter RCA species encode the *puf*M operon. In contrast, the deeply branching LUX cluster with streamlined genomes and a low G+C content and the COR cluster, in particular its subclusters 1 and 2 also exhibiting streamlined genomes and a low G+C content, were detected exclusively or predominantly from tropical to temperate regions (Fig. 6, Supplementary Fig. S8A). Only species COR-3-C, -5-E and -F3 were detected in all temperature regions and ocean basins (Supplementary Fig. S8). The great majority of the species of both clusters encode the *puf*LMX operon and those of the LUX cluster also the CBB cycle for CO_2_ fixation. Several species of the COR-6 subcluster, however, encode the PR gene (Supplementary Table S15). Hence, it appears that the deeply branching clusters evolved into two directions:

The TCR, AAPR and AAR clusters adapted more to temperate and cold environments and acquired the PR gene for binding complementary light energy. In contrast, the LUX cluster adapted to warmer regions and acquired the *puf*LMX operon for binding complementary light energy and most species even the CBB cycle to fix CO_2_. Future studies need to collect more data from ocean regions which were underrepresented in our dataset to further substantiate this observation. Most interestingly, this evolutionary divergence happened again in higher ranks of the phylogenetic tree of the *Roseobacter* group within the RCA and COR clusters: All species of the genus *Pseudoplanktomarina* and most species of the genus *Planktomarina* of the RCA cluster with typical features of genome streamlining, reduced genome size and G+C content relative to the most deeply branching RCA genus *Cand*. Paraplanktomarina (Table 1), acquired the PR gene. These species are distributed more in temperate and polar regions [7]. Three *Planktomarina* species, however with larger genomes and a higher G+C content than the most deeply branching genus of this cluster, acquired the *puf*M operon for AAP. Although the global biogeography of these species is not available in great detail it appears that these species occur predominantly in more nutrient-rich coastal areas of the temperate region [16,91,92]. In the COR cluster, subclusters 1 and 2 with pronounced genome streamlining features (Table 1), acquired the *puf*M operon and dwell predominantly in subtropical and tropical regions whereas several species of the COR-5 and -6 subclusters acquired the PR gene and were recruited and dwell predominantly in temperate to polar regions (Supplementary Fig. S8 and Table S11). It would be most interesting to elucidate the reasons for this dichotomic adaptation to acquiring complementary light energy occurring twice in the *Roseobacter* group. The lower energy cost for the biosynthesis but also the lower yield of energy bound by PR as compared to that of AAP and bacteriochlorophyll is known [93]. Here certainly other constraints including but not limited to temperature adaptation, nutrient and light availability play a role. Such information may be also important to better understand why the great majority of pelagic marine bacteria, the SAR11, SAR86, SAR116 and SAR202 clades, use PR to acquire complementary light energy. The fact that the LUX and COR clusters encode a much greater variety of genes to use organic phosphorous than the other pelagic clusters (Fig. 4A) appears to be another adaptation to the more or permanently stratified temperate and warm regions in which availability of inorganic phosphate is often extremely low [79].

Relationships between biogeographic patterns and genomic and functional features have also been studied for other prominent pelagic marine bacterial orders and/or families. Kent et al. [27,28] found distinct global biogeographic patterns of *Prochlorococcus* and *Synechococcus* sublineages in the tropical and subtropical oceans. These patterns were related to gene content and gene features associated to uptake and metabolism of nitrogen and phosphorous compounds and to light availability in near-surface waters and the deep chlorophyll maximum. Comprehensive studies have been carried out on the global biogeography of the SAR11 clade and subclades which showed detailed links between distribution patterns, ecotype speciation and functional features related to gene content, uptake and metabolism of organic and inorganic nutrients [20,30,94]. Roda-Garcia et al. [21] report distinct global distribution patterns of sublineages of the HGC and LGC subclades of the SAR116 clade, differing also greatly in their genome sizes and exhibiting some differences in the metabolism of organic sulfur compounds which may be related to biogeographic patterns. This study did not include the Arctic and Southern Oceans in which the SAR116 clade has been detected [95,96]. A global biogeography of the SAR86 clade and its four families, differing in genome size and functional features, has also been established recently. Representatives of all four families have been found in the major oceans, the Mediterranean and the Red Sea, however, with different relative abundances [31]. The SAR202 clade exhibits also a global biogeography [32]. The non-streamlined subgroups are predominantly dwelling in the meso- and bathypelagic whereas genome streamlined subgroups, evolved from different subgroups of the non-genome streamlined lineages, occur in the epipelagic of the different ocean basins in the Atlantic, Pacific and Indian Ocean [32]. Our investigation adds to these studies with a comprehensive analysis of the pelagic clusters of the *Roseobacter* group, providing detailed data on the genomic and functional features and global biogeography of these clusters, subclusters and species. It contributes to a much better understanding of how these clusters evolved and adapted to the different environmental conditions in the various ocean regions, predominantly in the epipelagic.

## Supplementary Information

## Acknowledgements

We want to thank all researchers and crew members of research vessels who collected the samples in the global oceans which were the basis for metagenomics and SAGs analyses and enabled us to recruit MAGs and SAGs of the *Roseobacter* group. We also want to thank Shinichi Sunagawa, Lucas Paoli and Rudi Amann for helpful discussions and assistance in collecting MAGs and SAGs from various databases.

## Authors’ contributions

YL carried out the analyses and wrote parts of the manuscript. MS and TB designed the study and MS wrote parts of the manuscript. All authors contributed to the interpretation and understanding of the findings and commented on the results and read the manuscript.

## Availability of data and materials

The genomes of 654 Roseobacter MAGs and SAGs analyzed in this study, along with their Prokka-predicted gene functions and the PR and *pufM* gene sequences used for phylogenetic reconstruction, are available on figshare (https://doi.org/10.6084/m9.figshare.29023568.v1)

## Declarations

Ethics approval and consent to participate Not applicable.

## Reference

1. Simon M, Scheuner C, Meier-Kolthoff JP, Brinkhoff T, Wagner-Döbler I, Ulbrich M, et al. Phylogenomics of Rhodobacteraceae reveals evolutionary adaptation to marine and non-marine habitats. ISME J. 2017;11:1483–99.

2. Liang KYH, Orata FD, Boucher YF, Case RJ. Roseobacters in a Sea of Poly- and Paraphyly: Whole Genome-Based Taxonomy of the Family Rhodobacteraceae and the Proposal for the Split of the “Roseobacter Clade” Into a Novel Family, Roseobacteraceae fam. nov. Front Microbiol. 2021;12:683109.

3. Giebel H-A, Kalhoefer D, Lemke A, Thole S, Gahl-Janssen R, Simon M, et al. Distribution of Roseobacter RCA and SAR11 lineages in the North Sea and characteristics of an abundant RCA isolate. ISME J. 2011;5:8–19.

4. Billerbeck S, Wemheuer B, Voget S, Poehlein A, Giebel H-A, Brinkhoff T, et al. Biogeography and environmental genomics of the Roseobacter-affiliated pelagic CHAB-I-5 lineage. Nat Microbiol. 2016;1:16063.

5. O’Brien J, McParland EL, Bramucci AR, Siboni N, Ostrowski M, Kahlke T, et al. Biogeographical and seasonal dynamics of the marine Roseobacter community and ecological links to DMSP-producing phytoplankton. ISME Commun. 2022;2:16.

6. Malmstrom R, Straza T, Cottrell M, Kirchman D. Diversity, abundance, and biomass production of bacterial groups in the western Arctic Ocean. Aquat Microb Ecol. 2007;47:45–55.

7. Liu Y, Brinkhoff T, Berger M, Poehlein A, Voget S, Paoli L, et al. Metagenome-assembled genomes reveal greatly expanded taxonomic and functional diversification of the abundant marine Roseobacter RCA cluster. Microbiome. 2023;11:265.

8. Segev E, Wyche TP, Kim KH, Petersen J, Ellebrandt C, Vlamakis H, et al. Dynamic metabolic exchange governs a marine algal-bacterial interaction. eLife. 2016;5:e17473.

9. Shibl AA, Isaac A, Ochsenkühn MA, Cárdenas A, Fei C, Behringer G, et al. Diatom modulation of select bacteria through use of two unique secondary metabolites. Proc Natl Acad Sci USA. 2020;117:27445–55.

10. Tran QD, Neu TR, Sultana S, Giebel H, Simon M, Billerbeck S. Distinct glycoconjugate cell surface structures make the pelagic diatom Thalassiosira rotula an attractive habitat for bacteria. J Phycol. 2023;59:309–22.

11. Selje N, Simon M, Brinkhoff T. A newly discovered Roseobacter cluster in temperate and polar oceans. Nature. 2004;427:445–8.

12. Moran MA, Durham BP. Sulfur metabolites in the pelagic ocean. Nat Rev Microbiol. 2019;17:665–78.

13. Swan BK, Tupper B, Sczyrba A, Lauro FM, Martinez-Garcia M, González JM, et al. Prevalent genome streamlining and latitudinal divergence of planktonic bacteria in the surface ocean. Proc National Acad Sci USA. 2013;110:11463–8.

14. Luo H, Moran MA. Evolutionary Ecology of the Marine Roseobacter Clade. Microbiol Mol Biol R. 2014;78:573–87.

15. Varaljay VA, Robidart J, Preston CM, Gifford SM, Durham BP, Burns AS, et al. Single-taxon field measurements of bacterial gene regulation controlling DMSP fate. ISME J. 2015;9:1677–86.

16. Voget S, Wemheuer B, Brinkhoff T, Vollmers J, Dietrich S, Giebel H-A, et al. Adaptation of an abundant Roseobacter RCA organism to pelagic systems revealed by genomic and transcriptomic analyses. ISME J. 2015;9:371–84.

17. Milke F, Meyerjürgens J, Simon M. Ecological mechanisms and current systems shape the modular structure of the global oceans’ prokaryotic seascape. Nat Commun. 2023;14:6141.

18. Luo H, Swan BK, Stepanauskas R, Hughes AL, Moran MA. Evolutionary analysis of a streamlined lineage of surface ocean Roseobacters. ISME J. 2014;8:1428–39.

19. Luo H, Csűros M, Hughes AL, Moran MA. Evolution of Divergent Life History Strategies in Marine Alphaproteobacteria. mBio. 2013;4:e00373–13.

20. Giovannoni SJ, Thrash JC, Temperton B. Implications of streamlining theory for microbial ecology. ISME J. 2014;8:1553–65.

21. Roda-Garcia JJ, Haro-Moreno JM, Huschet LA, Rodriguez-Valera F, López-Pérez M. Phylogenomics of SAR116 Clade Reveals Two Subclades with Different Evolutionary Trajectories and an Important Role in the Ocean Sulfur Cycle. Msystems. 2021;6:e00944–21.

22. Mende DR, Bryant JA, Aylward FO, Eppley JM, Nielsen T, Karl DM, et al. Environmental drivers of a microbial genomic transition zone in the ocean’s interior. Nat Microbiol. 2017;2:1367–73.

23. Graham ED, Heidelberg JF, Tully BJ. Potential for primary productivity in a globally-distributed bacterial phototroph. ISME J. 2018;12:1861–6.

24. Pachiadaki MG, Brown JM, Brown J, Bezuidt O, Berube PM, Biller SJ, et al. Charting the Complexity of the Marine Microbiome through Single-Cell Genomics. Cell. 2019;179:1623–1635.e11.

25. Dlugosch L, Poehlein A, Wemheuer B, Pfeiffer B, Badewien TH, Daniel R, et al. Significance of gene variants for the functional biogeography of the near-surface Atlantic Ocean microbiome. Nat Commun. 2022;13:456.

26. Biller SJ, Berube PM, Lindell D, Chisholm SW. Prochlorococcus: the structure and function of collective diversity. Nat Rev Microbiol. 2015;13:13–27.

27. Kent AG, Dupont CL, Yooseph S, Martiny AC. Global biogeography of Prochlorococcus genome diversity in the surface ocean. ISME J. 2016;10:1856–65.

28. Kent AG, Baer SE, Mouginot C, Huang JS, Larkin AA, Lomas MW, et al. Parallel phylogeography of Prochlorococcus and Synechococcus. ISME J. 2019;13:430–41.

29. Haro-Moreno JM, Rodriguez-Valera F, Rosselli R, Martinez-Hernandez F, Roda-Garcia JJ, Gomez ML, et al. Ecogenomics of the SAR11 clade. Environ Microbiol. 2020;22:1748–63.

30. Freel KC, Tucker SJ, Freel EB, Giovannoni SJ, Eren AM, Rappé MS. New isolate genomes and global marine metagenomes resolve ecologically relevant units of SAR11. bioRxiv. 2024.12.24.630191.

31. Ramfelt O, Freel KC, Tucker SJ, Nigro OD, Rappé MS. Isolate-anchored comparisons reveal evolutionary and functional differentiation across SAR86 marine bacteria. ISME J. 2024;18:wrae227.

32. He C, Gonsior M, Liu J, Jiao N, Chen F. Genome-streamlined SAR202 bacteria are widely present and active in the euphotic ocean. ISME J. 2025;19:wraf049.

33. Parks DH, Imelfort M, Skennerton CT, Hugenholtz P, Tyson GW. CheckM: assessing the quality of microbial genomes recovered from isolates, single cells, and metagenomes. Genome Res. 2015;25:1043–55.

34. Eren AM, Esen ÖC, Quince C, Vineis JH, Morrison HG, Sogin ML, et al. Anvi’o: an advanced analysis and visualization platform for ’omics data. PeerJ. 2015;3:e1319.

35. Olm MR, Brown CT, Brooks B, Banfield JF. dRep: a tool for fast and accurate genomic comparisons that enables improved genome recovery from metagenomes through de-replication. ISME J. 2017;11:2864–8.

36. Chaumeil P-A, Mussig AJ, Hugenholtz P, Parks DH. GTDB-Tk: a toolkit to classify genomes with the Genome Taxonomy Database. Bioinformatics. 2019;36:1925–7.

37. Minh BQ, Schmidt HA, Chernomor O, Schrempf D, Woodhams MD, Haeseler A von, et al. IQ-TREE 2: New models and efficient methods for phylogenetic inference in the genomic era. Mol Biol Evol. 2020;37:1530–4.

38. Letunic I, Bork P. Interactive Tree Of Life (iTOL) v5: an online tool for phylogenetic tree display and annotation. Nucleic Acids Res. 2021;49:W293–6.

39. Steinegger M, Söding J. MMseqs2 enables sensitive protein sequence searching for the analysis of massive data sets. Nat Biotechnol. 2017;35:1026–8.

40. Katoh K, Misawa K, Kuma K, Miyata T. MAFFT: a novel method for rapid multiple sequence alignment based on fast Fourier transform. Nucleic Acids Res. 2002;30:3059–66.

41. Capella-Gutiérrez S, Silla-Martínez JM, Gabaldón T. trimAl: a tool for automated alignment trimming in large-scale phylogenetic analyses. Bioinformatics. 2009;25:1972–3.

42. Parks DH, Rinke C, Chuvochina M, Chaumeil P-A, Woodcroft BJ, Evans PN, et al. Recovery of nearly 8,000 metagenome-assembled genomes substantially expands the tree of life. Nat Microbiol. 2017;2:1533–42.

43. Seemann T. Prokka: rapid prokaryotic genome annotation. Bioinformatics. 2014;30:2068–9.

44. Syberg-Olsen MJ, Garber AI, Keeling PJ, McCutcheon JP, Husnik F. Pseudofinder: Detection of Pseudogenes in Prokaryotic Genomes. Mol Biol Evol. 2022;39:msac153.

45. Aramaki T, Blanc-Mathieu R, Endo H, Ohkubo K, Kanehisa M, Goto S, et al. KofamKOALA: KEGG Ortholog assignment based on profile HMM and adaptive score threshold. Bioinformatics. 2020;36:2251–2.

46. Zhou Z, Tran PQ, Breister AM, Liu Y, Kieft K, Cowley ES, et al. METABOLIC: high-throughput profiling of microbial genomes for functional traits, metabolism, biogeochemistry, and community-scale functional networks. Microbiome. 2022;10:33.

47. Kanehisa M, Sato Y, Kawashima M, Furumichi M, Tanabe M. KEGG as a reference resource for gene and protein annotation. Nucleic Acids Res. 2016;44:D457–62.

48. Saier MH, Reddy VS, Moreno-Hagelsieb G, Hendargo KJ, Zhang Y, Iddamsetty V, et al. The Transporter Classification Database (TCDB): 2021 update. Nucleic Acids Res. 2020;49:D461–7.

49. Buchfink B, Reuter K, Drost H-G. Sensitive protein alignments at tree-of-life scale using DIAMOND. Nat Methods. 2021;18:366–8.

50. Paoli L, Ruscheweyh H-J, Forneris CC, Hubrich F, Kautsar S, Bhushan A, et al. Biosynthetic potential of the global ocean microbiome. Nature. 2022;1–8.

51. Nishimura Y, Yoshizawa S. The OceanDNA MAG catalog contains over 50,000 prokaryotic genomes originated from various marine environments. Sci Data. 2022;9:305.

52. Sánchez P, Coutinho FH, Sebastián M, Pernice MC, Rodríguez-Martínez R, Salazar G, et al. Marine picoplankton metagenomes and MAGs from eleven vertical profiles obtained by the Malaspina Expedition. Sci Data. 2024;11:154.

53. Cao S, Zhang W, Ding W, Wang M, Fan S, Yang B, et al. Structure and function of the Arctic and Antarctic marine microbiota as revealed by metagenomics. Microbiome. 2020;8:47.

54. Wood DE, Lu J, Langmead B. Improved metagenomic analysis with Kraken 2. Genome Biol. 2019;20:257.

55. Lu J, Breitwieser FP, Thielen P, Salzberg SL. Bracken: estimating species abundance in metagenomics data. PeerJ Comput Sci. 2017;3:e104.

56. Youngblut ND, Ley RE. Struo2: efficient metagenome profiling database construction for ever-expanding microbial genome datasets. PeerJ. 2021;9:e12198.

57. Durham BP, Grote J, Whittaker KA, Bender SJ, Luo H, Grim SL, et al. Draft genome sequence of marine alphaproteobacterial strain HIMB11, the first cultivated representative of a unique lineage within the Roseobacter clade possessing an unusually small genome. Stand Genom Sci. 2014;9:632–45.

58. Lanclos VC, Feng X, Cheng C, Yang M, Hider CJ, Coelho JT, et al. New isolates refine the ecophysiology of the Roseobacter CHAB-I-5 lineage. ISME Commun. 2025;ycaf068.

59. Zhang Z, Wu Z, Liu H, Yang M, Wang R, Zhao Y, et al. Genomic analysis and characterization of phages infecting the marine Roseobacter CHAB-I-5 lineage reveal a globally distributed and abundant phage genus. Front Microbiol. 2023;14:1164101.

60. Feng X, Chu X, Qian Y, Henson MW, Lanclos VC, Qin F, et al. Mechanisms driving genome reduction of a novel Roseobacter lineage. ISME J. 2021;1–11.

61. Giovannoni SJ. SAR11 Bacteria: The Most Abundant Plankton in the Oceans. Annu Rev Mar Sci. 2017;9:231–55.

62. Yang Y, Wang P, Qaidi SE, Hardwidge PR, Huang J, Zhu G. Loss to gain: pseudogenes in microorganisms, focusing on eubacteria, and their biological significance. Appl Microbiol Biotechnol. 2024;108:328.

63. Liu Y, Harrison PM, Kunin V, Gerstein M. Comprehensive analysis of pseudogenes in prokaryotes: widespread gene decay and failure of putative horizontally transferred genes. Genome Biol. 2004;5:R64–R64.

64. Pinhassi J, DeLong EF, Béjà O, González JM, Pedrós-Alió C. Marine Bacterial and Archaeal Ion-Pumping Rhodopsins: Genetic Diversity, Physiology, and Ecology. Microbiol Mol Biology Rev Mmbr. 2016;80:929–54.

65. Vollmers J, Voget S, Dietrich S, Gollnow K, Smits M, Meyer K, et al. Poles Apart: Arctic and Antarctic Octadecabacter strains Share High Genome Plasticity and a New Type of Xanthorhodopsin. PLoS ONE. 2013;8:e63422.

66. Newton RJ, Griffin LE, Bowles KM, Meile C, Gifford S, Givens CE, et al. Genome characteristics of a generalist marine bacterial lineage. ISME J. 2010;4:784–98.

67. Giebel H-A, Wolterink M, Brinkhoff T, Simon M. Complementary energy acquisition via aerobic anoxygenic photosynthesis and carbon monoxide oxidation by Planktomarina temperata of the Roseobacter group. Fems Microbiol Ecol. 2019;95.

68. Pitcher RS, Watmough NJ. The bacterial cytochrome cbb3 oxidases. Biochim Biophys Acta (BBA) - Bioenerg. 2004;1655:388–99.

69. Borisov VB, Gennis RB, Hemp J, Verkhovsky MI. The cytochrome bd respiratory oxygen reductases. Biochim Biophys Acta (BBA) - Bioenerg. 2011;1807:1398–413.

70. Anantharaman K, Hausmann B, Jungbluth SP, Kantor RS, Lavy A, Warren LA, et al. Expanded diversity of microbial groups that shape the dissimilatory sulfur cycle. ISME J. 2018;12:1715–28.

71. Klingner A, Bartsch A, Dogs M, Wagner-Döbler I, Jahn D, Simon M, et al. Large-Scale 13C Flux Profiling Reveals Conservation of the Entner-Doudoroff Pathway as a Glycolytic Strategy among Marine Bacteria That Use Glucose. Appl Environ Microbiol. 2015;81:2408–22.

72. Palovaara J, Akram N, Baltar F, Bunse C, Forsberg J, Pedrós-Alió C, et al. Stimulation of growth by proteorhodopsin phototrophy involves regulation of central metabolic pathways in marine planktonic bacteria. Proc Natl Acad Sci USA. 2014;111:E3650–8.

73. Schneider K, Peyraud R, Kiefer P, Christen P, Delmotte N, Massou S, et al. The Ethylmalonyl-CoA Pathway Is Used in Place of the Glyoxylate Cycle by Methylobacterium extorquens AM1 during Growth on Acetate. J Biol Chem. 2012;287:757–66.

74. Lidbury ID, Murrell JC, Chen Y. Trimethylamine and trimethylamine N-oxide are supplementary energy sources for a marine heterotrophic bacterium: implications for marine carbon and nitrogen cycling. ISME J. 2015;9:760–9.

75. Lidbury I, Mausz MA, Scanlan DJ, Chen Y. Identification of dimethylamine monooxygenase in marine bacteria reveals a metabolic bottleneck in the methylated amine degradation pathway. ISME J. 2017;11:1592–601.

76. Grevesse T, Guéguen C, Onana VE, Walsh DA. Degradation pathways for organic matter of terrestrial origin are widespread and expressed in Arctic Ocean microbiomes. Microbiome. 2022;10:237.

77. Stasi R, Neves HI, Spira B. Phosphate uptake by the phosphonate transport system PhnCDE. BMC Microbiol. 2019;19:79.

78. Koedooder C, Zhang F, Wang S, Basu S, Haley ST, Tolic N, et al. Taxonomic distribution of metabolic functions in bacteria associated with Trichodesmium consortia. mSystems. 2023;8:e00742–23.

79. Karl DM. Microbially Mediated Transformations of Phosphorus in the Sea: New Views of an Old Cycle. Annu Rev Mar Sci. 2014;6:279–337.

80. Hagström Å, Zweifel UL, Sundh J, Osbeck CMG, Bunse C, Sjöstedt J, et al. Composition and Seasonality of Membrane Transporters in Marine Picoplankton. Front Microbiol. 2021;12:714732.

81. Clifton BE, Alcolombri U, Uechi G-I, Jackson CJ, Laurino P. The ultra-high affinity transport proteins of ubiquitous marine bacteria. Nature. 2024;634:721–8.

82. Ahmed MA, Campbell BJ. Genome-resolved adaptation strategies of Rhodobacterales to changing conditions in the Chesapeake and Delaware Bays. Appl Environ Microbiol. 2025;91:e02357–24.

83. Tang K, Jiao N, Liu K, Zhang Y, Li S. Distribution and Functions of TonB-Dependent Transporters in Marine Bacteria and Environments: Implications for Dissolved Organic Matter Utilization. PLoS ONE. 2012;7:e41204.

84. Zhao Z, Amano C, Reinthaler T, Orellana MV, Herndl GJ. Substrate uptake patterns shape niche separation in marine prokaryotic microbiome. Sci Adv. 2024;10:eadn5143.

85. Morris RM, Frazar CD, Carlson CA. Basin-scale patterns in the abundance of SAR11 subclades, marine Actinobacteria (OM1), members of the Roseobacter clade and OCS116 in the South Atlantic. Environ Microbiol. 2012;14:1133–44.

86. Wilkins D, Lauro FM, Williams TJ, Demaere MZ, Brown MV, Hoffman JM, et al. Biogeographic partitioning of Southern Ocean microorganisms revealed by metagenomics. Environ Microbiol. 2013;15:1318–33.

87. Tada Y, Shiozaki T, Ogawa H, Suzuki K. Basin-scale distribution of prokaryotic phylotypes in the epipelagic layer of the Central South Pacific Ocean during austral summer. J Oceanogr. 2017;73:145–58.

88. Bakenhus I, Dlugosch L, Giebel H, Beardsley C, Simon M, Wietz M. Distinct biogeographic patterns of bacterioplankton composition and single-cell activity between the subtropics and Antarctica. Environ Microbiol. 2018;20:3100–8.

89. Reintjes G, Tegetmeyer HE, Bürgisser M, Orlić S, Tews I, Zubkov M, et al. On-Site Analysis of Bacterial Communities of the Ultraoligotrophic South Pacific Gyre. Appl Environ Microbiol. 2019;85:e00184–19.

90. Salazar G, Cornejo-Castillo FM, Benítez-Barrios V, Fraile-Nuez E, Álvarez-Salgado XA, Duarte CM, et al. Global diversity and biogeography of deep-sea pelagic prokaryotes. ISME J. 2016;10:596–608.

91. Mayali X, Franks PJS, Azam F. Cultivation and Ecosystem Role of a Marine Roseobacter Clade-Affiliated Cluster Bacterium. Appl Environ Microb. 2008;74:2595–603.

92. Giebel H-A, Kalhoefer D, Gahl-Janssen R, Choo Y-J, Lee K, Cho J-C, et al. Planktomarina temperata gen. nov., sp. nov., belonging to the globally distributed RCA cluster of the marine Roseobacter clade, isolated from the German Wadden Sea. Int J Syst Evol Micr. 2013;63:4207–17.

93. Kirchman DL, Hanson TE. Bioenergetics of photoheterotrophic bacteria in the oceans. Env Microbiol Rep. 2013;5:188–99.

94. Delmont TO, Kiefl E, Kilinc O, Esen OC, Uysal I, Rappé MS, et al. Single-amino acid variants reveal evolutionary processes that shape the biogeography of a global SAR11 subclade. eLife. 2019;8:e46497.

95. Wietz M, Bienhold C, Metfies K, Torres-Valdés S, Appen W-J von, Salter I, et al. The polar night shift: seasonal dynamics and drivers of Arctic Ocean microbiomes revealed by autonomous sampling. ISME Commun. 2021;1:76.

96. West NJ, Landa M, Obernosterer I. Differential association of key bacterial groups with diatoms and Phaeocystis spp. during spring blooms in the Southern Ocean. MicrobiologyOpen. 2024;13:e1428.

