## Supplementary Figures for "Ecogenomics and functional biogeography of the *Roseobacter* group in the global oceans based on 653 MAGs and SAGs"

*Authors for correspondence:

**Supplementary Figures**

**Figure S1. Close-up view of the deeply branching pelagic clusters of the *Roseobacter* group**. MAGs, SAGs and genome HTC2255 of the TCR, AAPR, AAR and LUX clusters are shown together with the deeply branching genera *Pontivivens*, *Albimonas*, *Neptunicoccus* and *Paramylibacter*.

**Figure S2. Distribution of the genome size and G+C content of MAGs, SAGs and genomes of the pelagic clusters and of isolates of the *Roseobacter* group**. 154 representative genomes of MAGs, SAGS and genomes of the pelagic clusters and 344 reference genomes of isolates are included in these analyses. (A) BAR and density plot of the genome size. (B) BAR and density plot of the G+C content.

**Figure S3. Phylogeny and ANI of species of the AAR, AAPR, TCR, COR and LUX clusters.** (A) Phylogeny of the MAGs, SAGs and genomes of all species of the AAPR, AAR, TCR, COR and LUX cluster, species identification and recruitment region or latitudinal range (right of the tree in parenthesis). Each species is color coded. (B) ANI of the species of the AAPR, AAR, TCR, COR and LUX cluster.

**Figure S4. Close-up view of the phylogeny of the COR cluster and neighboring taxa.** *Pseudaestuariivita* and *Marinovum* are the most closely related genera to the COR cluster.

**Figure S5. Phylogenetic trees of *puf*M and rhodopsin genes of Bacteria.** (A) Phylogenetic tree of the rhodopsin (xanthorhodopsin, proteorhodopsin) gene based on 207 PR amino acid sequences from our dataset and 514 reference sequences from the UniProt database. Light absorbing properties are indicted in blue and green dots. Genome classification is indicated by different colors in the inner semicircle, with genomes from our dataset further highlighted by distinct colored backgrounds. Rhodopsin types are indicated in the outer semicircle. (B) The *puf*M phylogenetic tree was constructed from 100 *puf*M amino acid sequences from our dataset and 348 reference sequences of the bacterial phylum. Three *puf* operon types (*puf*LMX, *puf*LMC and *puf*LM) found in *Roseobacter* genomes analyzed in this study are indicated by distinct color dots. Seven genomes from the COR cluster contain both *puf*LMX and *puf*LMC operons. The outer and inner semicircles represent genome classifications at the class, genus or cluster level, respectively. The genomes from the our dataset are further highlighted by distinct colored backgrounds.

**Figure S6. Transporter distribution in species of AAR, AAR, RCA, TCA, COR and LUX clusters.** (A, C) Substrate-binding protein (SBP)–dependent and ABC transporters in different species (% of total genes). (B, D) SBP-dependent and ABC transporters per megabase (Mbp) in different species. Only species with more than four genomes are shown. Significant differences ($p < 0.05$) among different species are denoted by letters a to l.

**Figure S7. Relative abundance of the *Roseobacter* group in the epi- (A) meso- (B) and bathypelagic (C) as percent of total prokaryotes in the global oceans.** For details on the different ocean regions and temperature zones see Supplementary Tables S36 to S38.

**Figure S8. Relative abundance of different species in AAPR, AAR, RCA, TCR, COR, and LUX clusters in epipelagic waters across different temperature zones (A) and ocean regions (B).** Species abundance in clusters was measured based on the percentage of read recruitment using Kraken2 and Bracken. Only species recruited in more than five metagenomic samples are shown. Circle size indicates the relative abundance of each species. N and S represent the Northern and Southern Hemispheres, respectively. For details on the different clusters and genera see Supplementary Tables S39 to S44.

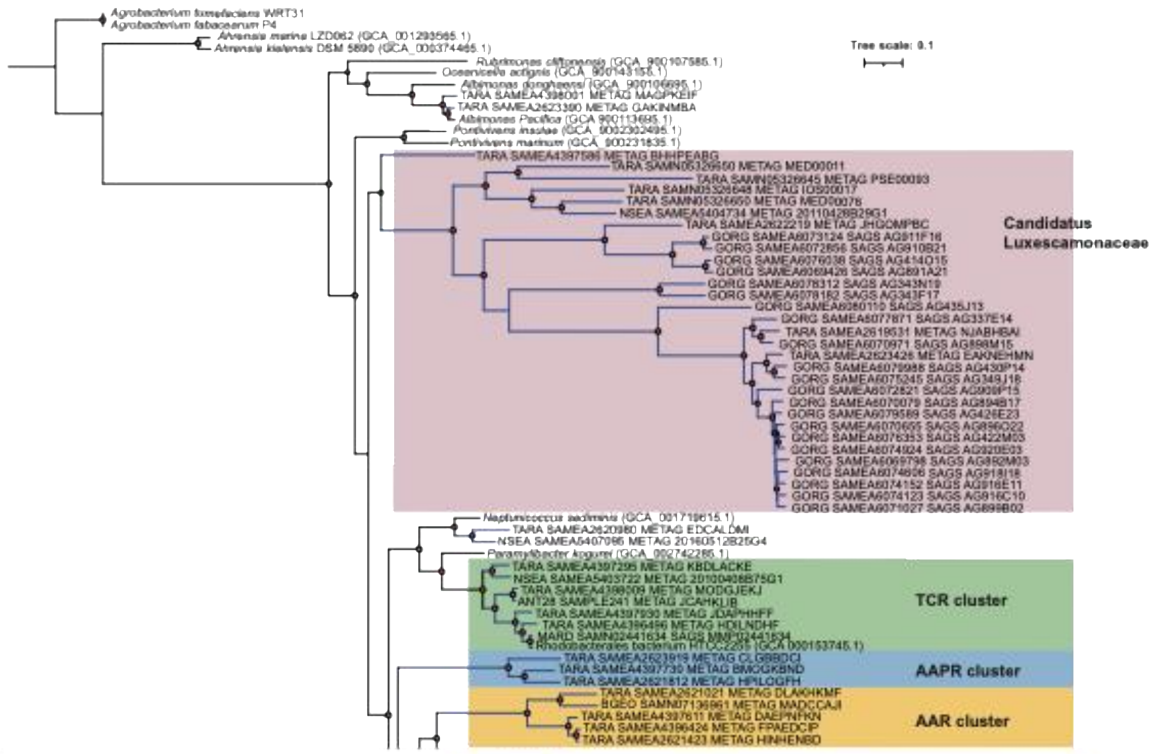


Figure S1

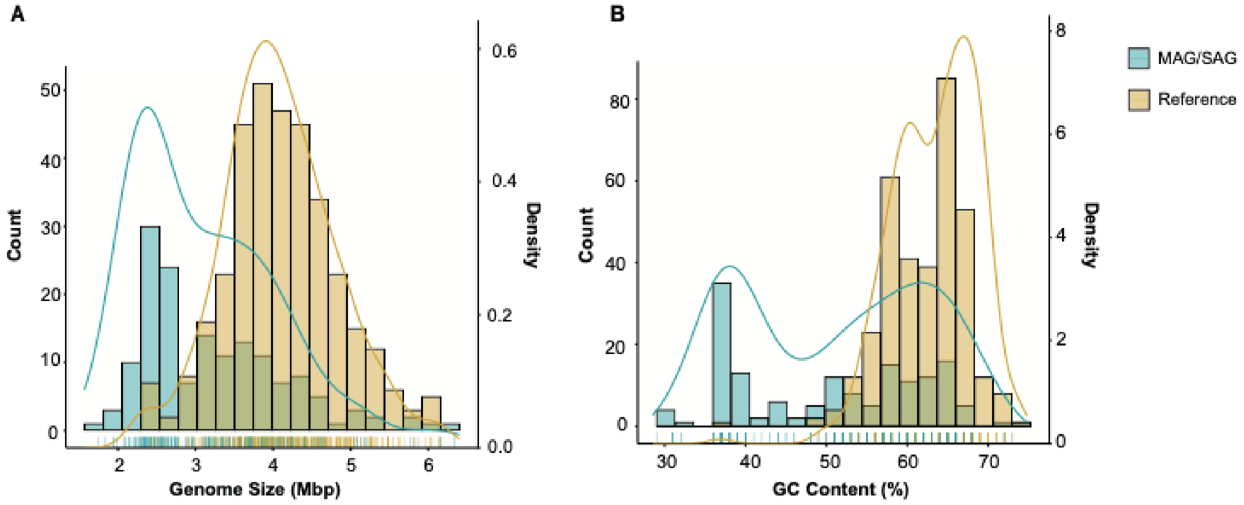


Figure S2

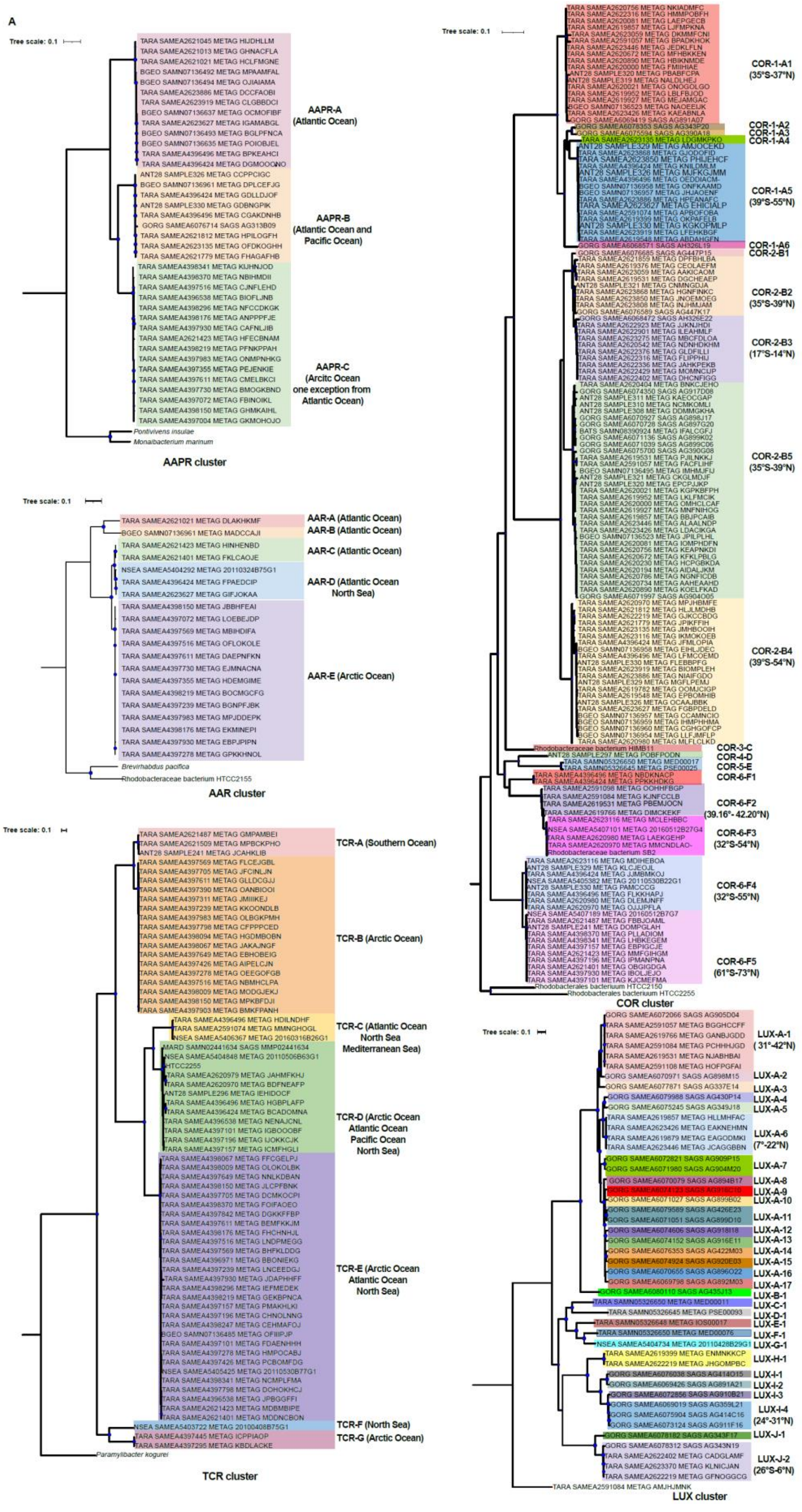
A
Tree scale: 0.1
AAPR-A (Atlantic Ocean)
AAPR-B (Atlantic Ocean and Pacific Ocean)
AAPR-C (Arcitc Ocean one exception from Atlantic Ocean)
Pontivivens insulae
Monaibacterium marinum
AAPR cluster
AAR-A (Atlantic Ocean)
AAR-B (Atlantic Ocean)
AAR-C (Atlantic Ocean)
AAR-D (Atlantic Ocean North Sea)
AAR-E (Arctic Ocean)
Brevirhabdus pacifica
Rhodobacteraceae bacterium HTCC2155
AAR cluster
TCR-A (Southern Ocean)
TCR-B (Arctic Ocean)
TCR-C (Atlantic Ocean North Sea Mediterranean Sea)
TCR-D (Arctic Ocean Atlantic Ocean Pacific Ocean North Sea)
TCR-E (Arctic Ocean Atlantic Ocean North Sea)
TCR-F (North Sea)
TCR-G (Arctic Ocean)
Paramylibacter kogurei
TCR cluster
COR-1-A1 (35°S-37°N)
COR-1-A2
COR-1-A3
COR-1-A4
COR-1-A5 (39°S-55°N)
COR-1-A6
COR-2-B1
COR-2-B2 (35°S-39°N)
COR-2-B3 (17°S-14°N)
COR-2-B5 (35°S-39°N)
COR-2-B4 (39°S-54°N)
COR-3-C
COR-4-D
COR-5-E
COR-6-F1
COR-6-F2 (39.16°- 42.20°N)
COR-6-F3 (32°S-54°N)
COR-6-F4 (32°S-55°N)
COR-6-F5 (61°S-73°N)
Rhodobacterales bacterium HTCC2150
Rhodobacterales bacterium HTCC2255
COR cluster
LUX-A-1 ( 31°-42°N)
LUX-A-2
LUX-A-3
LUX-A-4
LUX-A-5
LUX-A-6 (7°-22°N)
LUX-A-7
LUX-A-8
LUX-A-9
LUX-A-10
LUX-A-11
LUX-A-12
LUX-A-13
LUX-A-14
LUX-A-15
LUX-A-16
LUX-A-17
LUX-B-1
LUX-C-1
LUX-D-1
LUX-E-1
LUX-F-1
LUX-G-1
LUX-H-1
LUX-I-1
LUX-I-2
LUX-I-3
LUX-I-4 (24°-31°N)
LUX-J-1
LUX-J-2 (26°S-6°N)
LUX cluster

Figure S3A

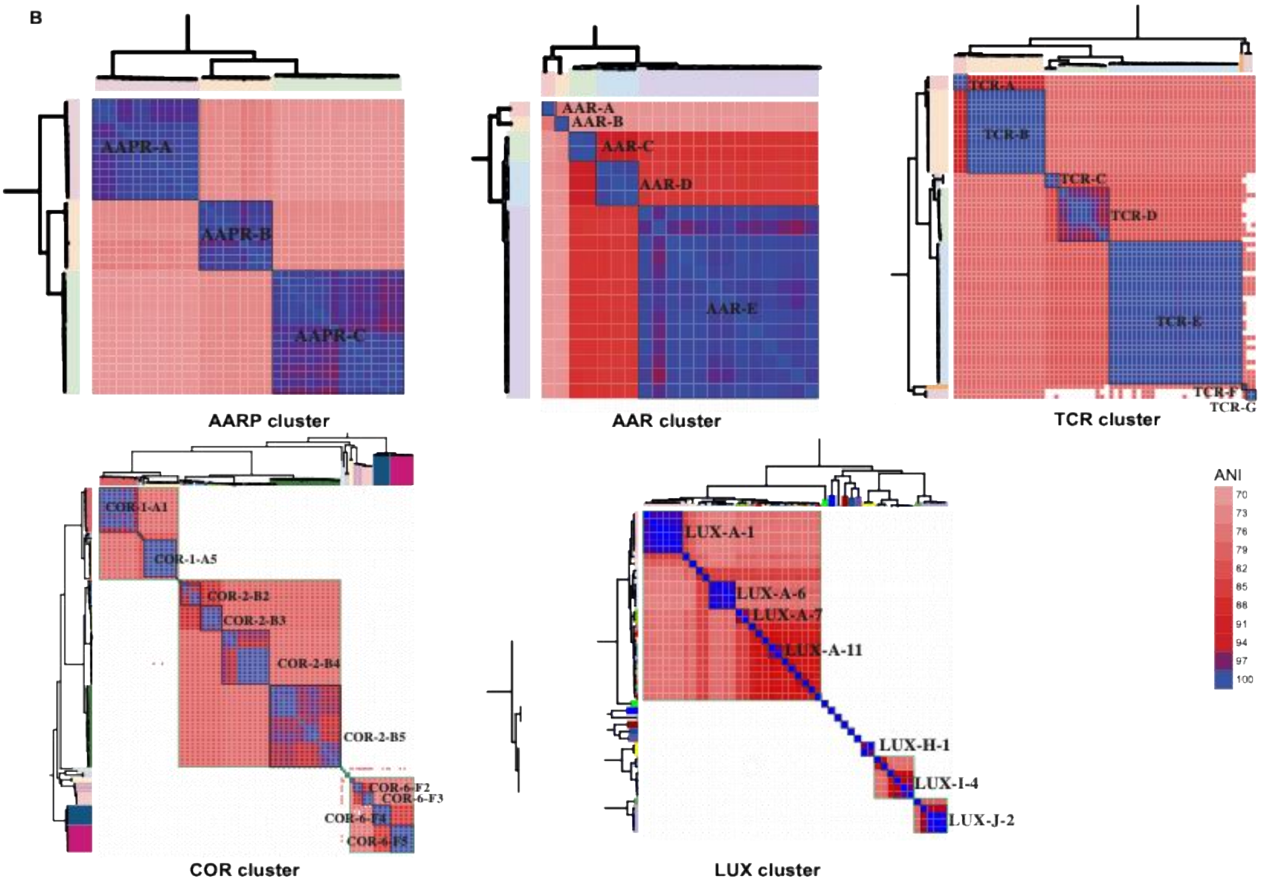
B
AAPR-A
AAPR-B
AAPR-C
AARP cluster
AAR-A
AAR-B
AAR-C
AAR-D
AAR-E
AAR cluster
TCR-A
TCR-B
TCR-C
TCR-D
TCR-E
TCR-F
TCR-G
TCR cluster
COR-1-A1
COR-1-A5
COR-2-B2
COR-2-B3
COR-2-B4
COR-2-B5
COR-6-F2
COR-6-F3
COR-6-F4
COR-6-F5
COR cluster
LUX-A-1
LUX-A-6
LUX-A-7
LUX-A-11
LUX-H-1
LUX-1-4
LUX-J-2
LUX cluster
ANI
70
73
76
79
82
85
88
91
94
97
100

Figure S3B

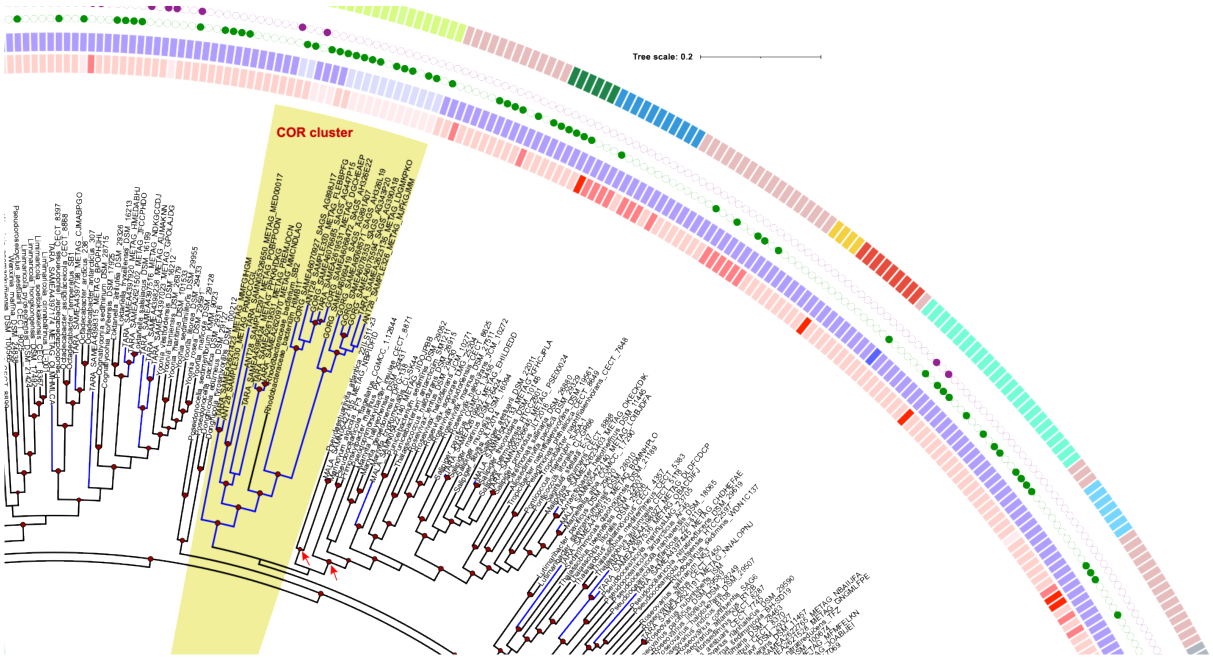


Figure S4

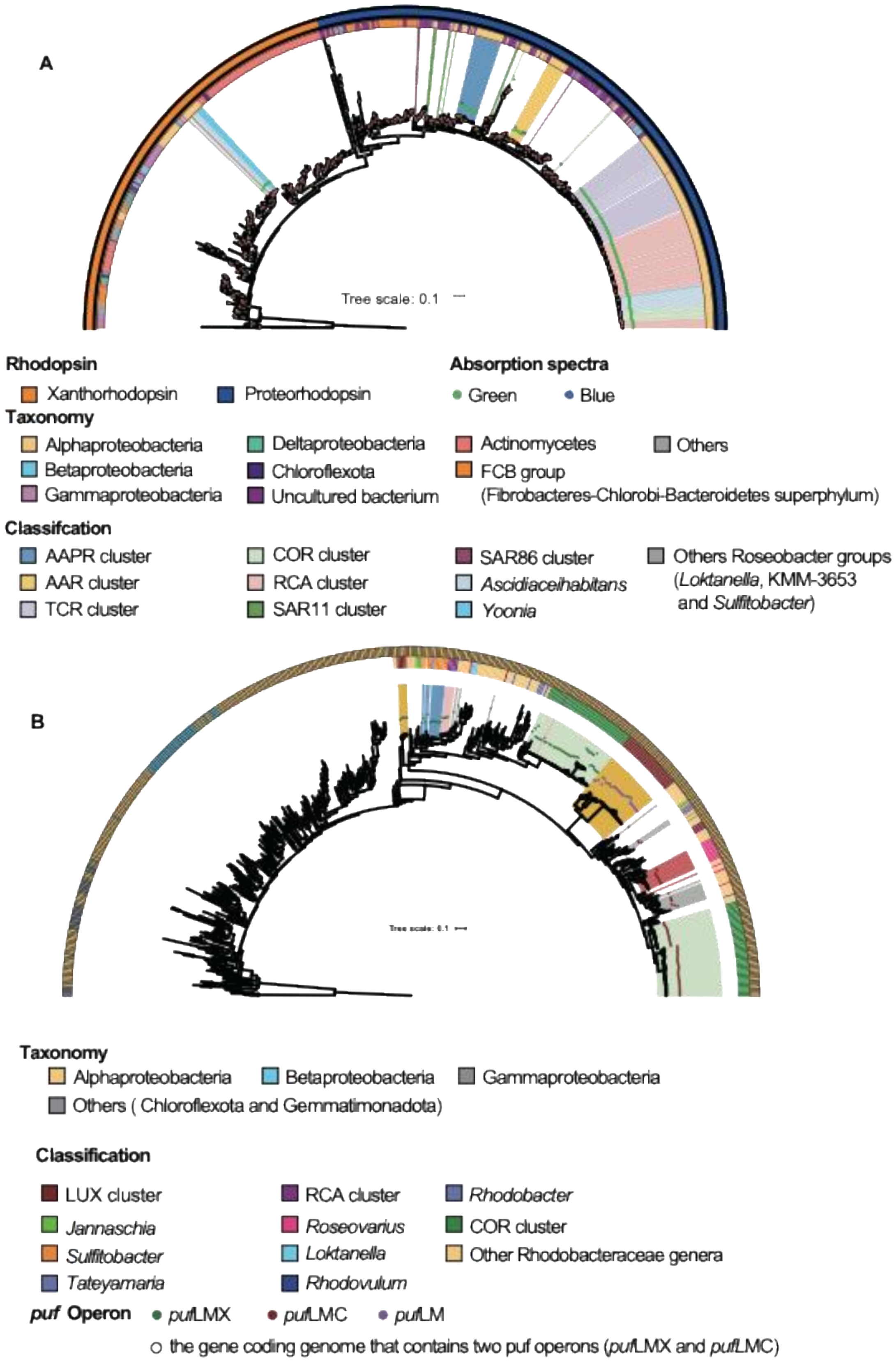
A
Tree scale: 0.1
Rhodopsin
Xanthorhodopsin
Proteorhodopsin
Absorption spectra
Green
Blue
Taxonomy
Alphaproteobacteria
Betaproteobacteria
Gammaproteobacteria
Deltaproteobacteria
Chloroflexota
Uncultured bacterium
Actinomycetes
FCB group
(Fibrobacteres-Chlorobi-Bacteroidetes superphylum)
Others
Classifcation
AAPR cluster
AAR cluster
TCR cluster
COR cluster
RCA cluster
SAR11 cluster
SAR86 cluster
Ascidiaceihabitans
Yoonia
Others Roseobacter groups
(Loktanella, KMM-3653
and Sulfitobacter)
B
Tree scale: 0.1
Taxonomy
Alphaproteobacteria
Betaproteobacteria
Gammaproteobacteria
Others ( Chloroflexota and Gemmatimonadota)
Classification
LUX cluster
Jannaschia
Sulfitobacter
Tateyamaria
RCA cluster
Roseovarius
Loktanella
Rhodovulum
Rhodobacter
COR cluster
Other Rhodobacteraceae genera
puf Operon
pufLMX
pufLMC
pufLM
the gene coding genome that contains two puf operons (pufLMX and pufLMC)

Figure S5

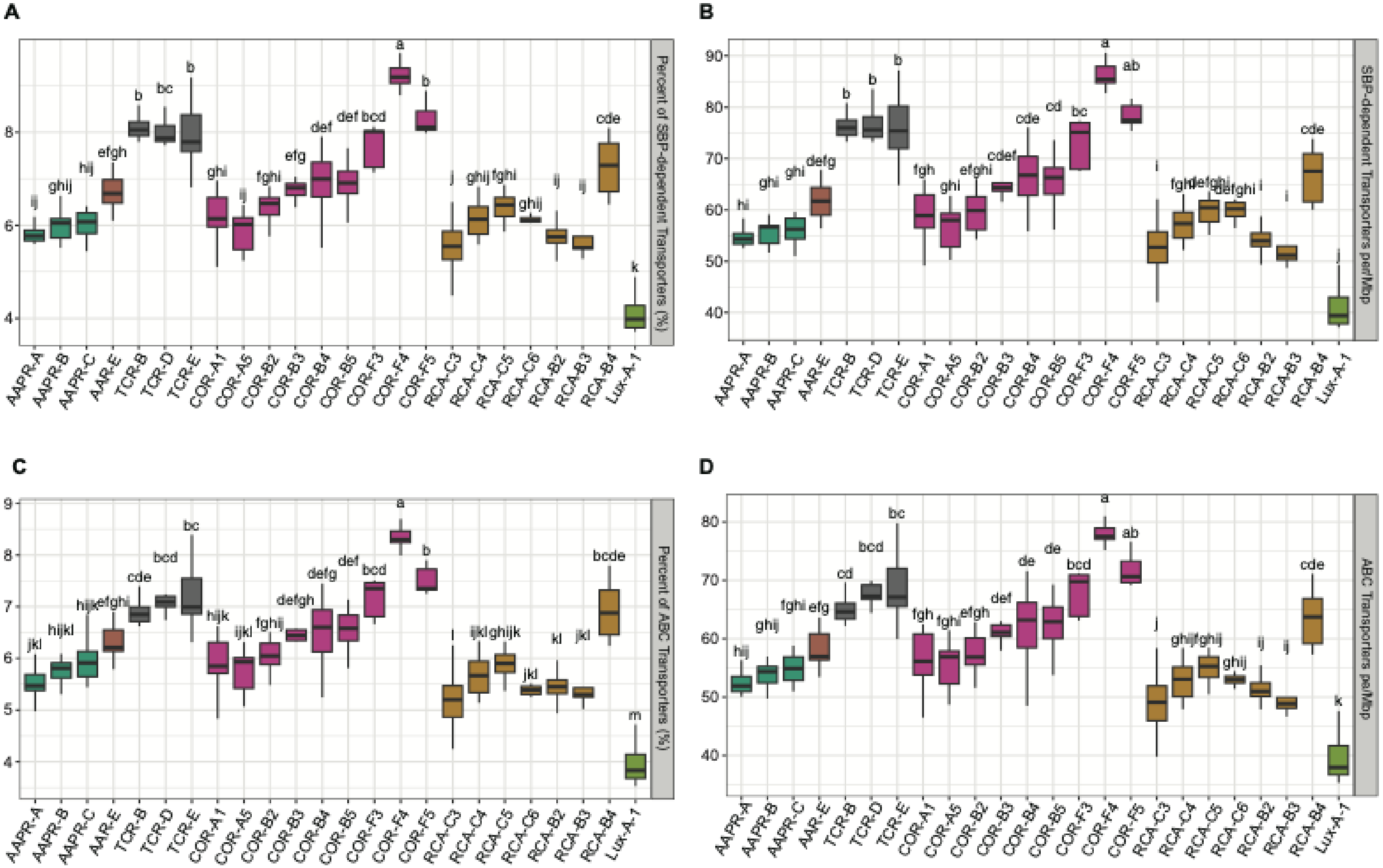
A
B
C
D
Percent of SBP-dependent Transporters (%)
SBP-dependent Transporters per Mbp
Percent of ABC Transporters (%)
ABC Transporters per Mbp
AAPR-A
AAPR-B
AAPR-C
AAR-E
TCR-B
TCR-D
TCR-E
COR-A1
COR-A5
COR-B2
COR-B3
COR-B4
COR-B5
COR-F3
COR-F4
COR-F5
RCA-C3
RCA-C4
RCA-C5
RCA-C6
RCA-B2
RCA-B3
RCA-B4
Lux-A-1

Figure S6

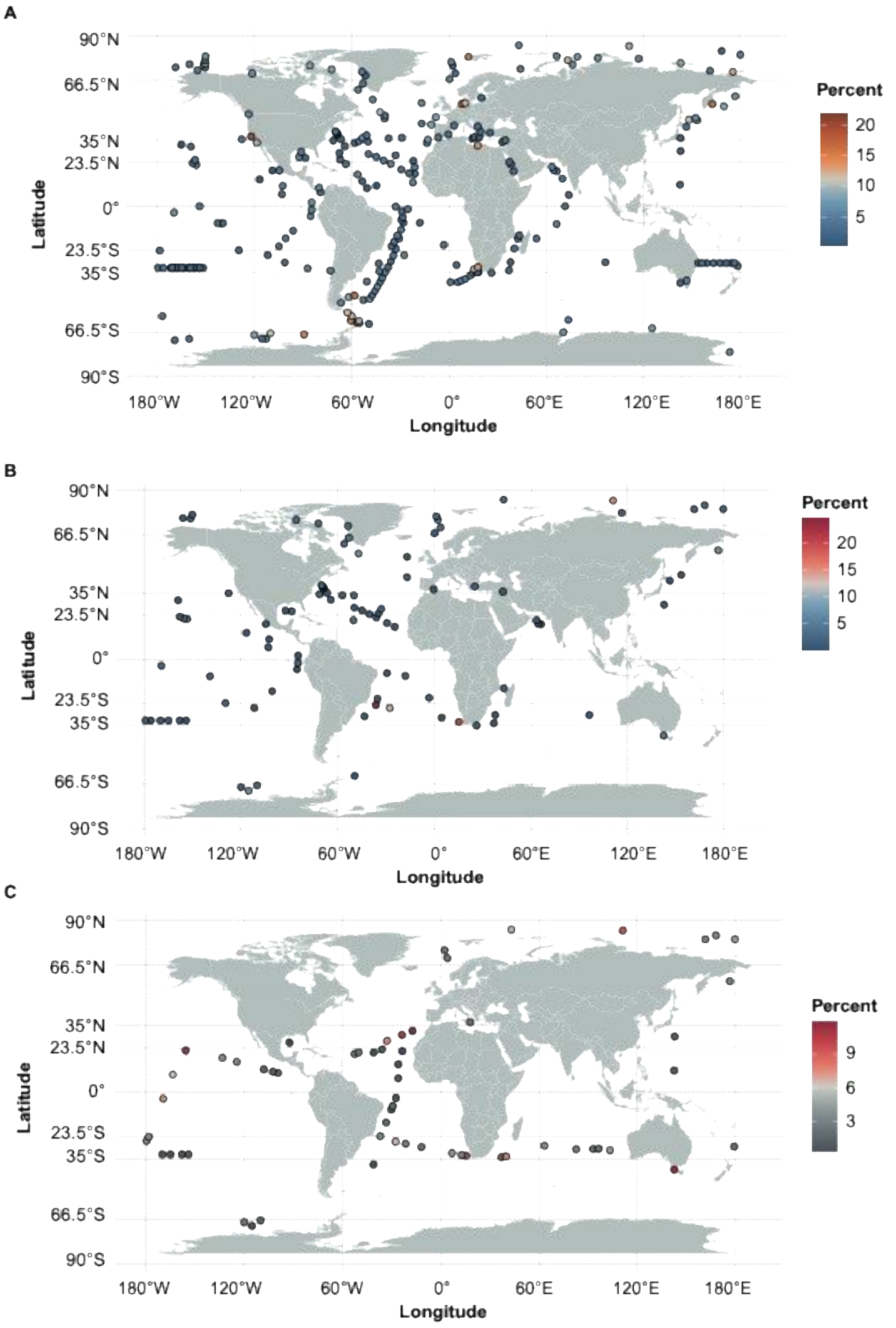
A
90°N
66.5°N
35°N
23.5°N
0°
23.5°S
35°S
66.5°S
90°S
Latitude
180°W
120°W
60°W
0°
60°E
120°E
180°E
Longitude
Percent
20
15
10
5
B
90°N
66.5°N
35°N
23.5°N
0°
23.5°S
35°S
66.5°S
90°S
Latitude
180°W
120°W
60°W
0°
60°E
120°E
180°E
Longitude
Percent
20
15
10
5
C
90°N
66.5°N
35°N
23.5°N
0°
23.5°S
35°S
66.5°S
90°S
Latitude
180°W
120°W
60°W
0°
60°E
120°E
180°E
Longitude
Percent
9
6
3

Figure S7

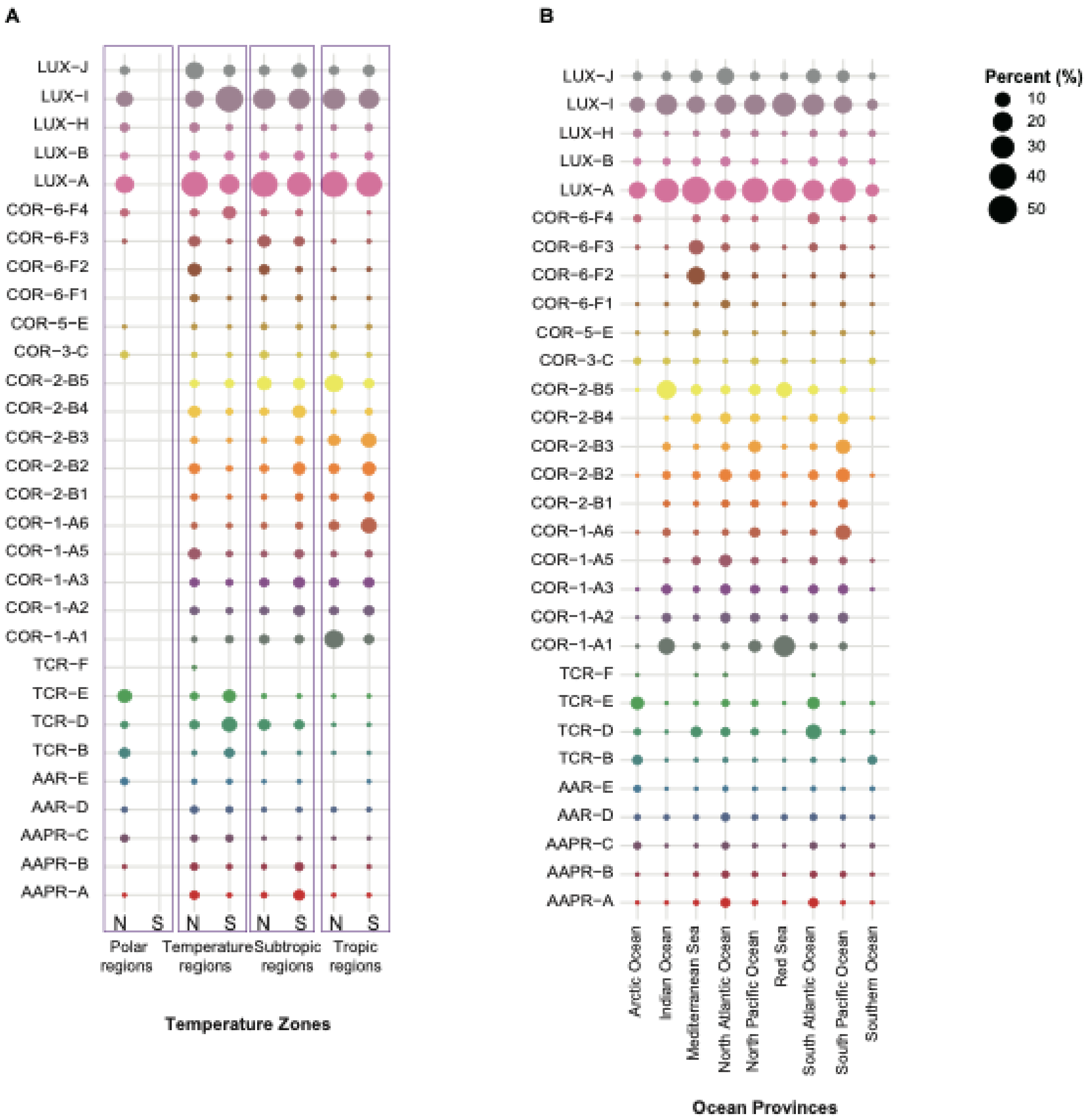
A
B
LUX-J
LUX-I
LUX-H
LUX-B
LUX-A
COR-6-F4
COR-6-F3
COR-6-F2
COR-6-F1
COR-5-E
COR-3-C
COR-2-B5
COR-2-B4
COR-2-B3
COR-2-B2
COR-2-B1
COR-1-A6
COR-1-A5
COR-1-A3
COR-1-A2
COR-1-A1
TCR-F
TCR-E
TCR-D
TCR-B
AAR-E
AAR-D
AAPR-C
AAPR-B
AAPR-A
N S N S N S N S
Polar regions
Temperature regions
Subtropic regions
Tropic regions
Temperature Zones
Arctic Ocean
Indian Ocean
Mediterranean Sea
North Atlantic Ocean
North Pacific Ocean
Red Sea
South Atlantic Ocean
South Pacific Ocean
Southern Ocean
Ocean Provinces
Percent (%)
10
20
30
40
50

Figure S8
