## Supplementary Text for "Ecogenomics and functional biogeography of the *Roseobacter* group in the global oceans based on 653 MAGs and SAGs"

*Authors for correspondence:

### Supplementary Text

#### Phosphorous acquisition by the pelagic *Roseobacter* clusters

Phosphorous acquisition is variable among the six clusters and less versatile than in most other genera of the *Roseobacter* group (Fig. 4B, Supplementary Table S24). In several MAGs of each species of the six clusters only very few or no phosphorous-related genes were detected, possibly an effect of the incomplete genomes. In these cases, we considered a gene present in a species when this gene was detected in at least one MAG of a given species. Species A of the

AAPR cluster encodes the high affinity phosphate transporter (*pst*ABCS) whereas species C and B the low affinity phosphate transporter (*pit*A), but no other gene related to phosphorous uptake (Supplementary Table S24). Species A to D of cluster AAR also encode either the low or high affinity phosphate transporter but in addition also the 5'-nucleotidase whereas several MAGs of species E encode only the 5'-nucleotidase and no phosphate transporter (Fig. 5, Supplementary Table S24). All except species C of the TCR cluster encode either the low or high affinity phosphate transporter or both and in addition a phosphonate transporter (*phn*CDE) even though no gene encoding the C-P lyase was detected. It has been shown, though, that the phosphonate transporter PhnCDE can also transport phosphate [1] making it likely that this is also the case for species of the pelagic *Roseobacter* clusters and other pelagic marine bacteria. The genus *Pseudoplanktomarina* and *Cand*. Paraplanktomarina of the RCA cluster only encode the high affinity phosphate transporter but in addition the 5'-nucleotidase whereas most species of the genus *Planktomarina* also encode the low affinity phosphate transporter and a few species the phosphonate transporter as well (Fig. 5, Supplementary Table S24). The great majority of MAGs of subclusters COR-1 and COR-2 lack the high and low affinity phosphate transporter and the subclusters COR-3 to COR-6 lack the low affinity phosphate transporter (Supplementary Table S24). However, most species of these subclusters encode alkaline phosphatase, the phosphonoacetate hydrolase and the phosphonate transporter and quite a few also the C-P lyase. Hence species of the COR cluster are able to exploit a variety of organic phosphorous compounds, an important resource when inorganic phosphate is severely limiting or unavailable [2,3]. Species of the LUX cluster are also predicted to be quite versatile in exploiting organic phosphorous compounds as they encode several genes catalyzing the cleavage of organic phosphorous bonds (Fig. 4B, Supplementary Table S24).

### Biogeography of single species and subclusters in the epipelagic

Looking into the diversity within the AAR cluster species D was detected in all ocean regions whereas species E was only detected in the Arctic and Atlantic Ocean (Supplementary Fig. S8B). Unfortunately, species AAR-A, -B and -C could not be resolved sufficiently in our kraken2 analysis, i.e. detected in < 5 metagenomes, and thus could not be considered in the biogeographic distribution. Regarding the AAPR cluster, species C was not detected in the Mediterranean Sea and the South Pacific Ocean. Species TCR-B was the only species of this cluster found in the Southern Ocean but not detected in the Pacific Ocean and the Mediterranean Sea. Species TCR-A, -C and -G did not pass the criteria of our kraken2 analysis and were not considered in the biogeographic analysis. All species of the genome streamlined subclusters COR-1 and COR-2 were detected only in the non-polar regions whereas species COR-6-F4 with the largest genome and the highest G+C content of the COR cluster (Table 1) was detected also in the polar regions of both hemispheres but not in the Indian Ocean and Red Sea (Supplementary Fig. S8B). Other species of subclusters COR-6 and COR-5 were not either detected in these warm regions. A recent study on the biogeography of two isolates affiliated to *Cand*. Thalassiovivens spotiae [4], closely related to isolate SB2 of cluster COR-6-F3 [5], reported an even broader occurrence than we found including also the polar regions of both hemispheres, the Indian Ocean and Red Sea. A detailed analysis of the biogeography of the different RCA species with preferences to the temperate and polar regions of both hemispheres has been published elsewhere [6].

**References**


1. Stasi R, Neves HI, Spira B. Phosphate uptake by the phosphonate transport system PhnCDE. BMC Microbiol. 2019;19:79.

2. Koedooder C, Zhang F, Wang S, Basu S, Haley ST, Tolic N, et al. Taxonomic distribution of metabolic functions in bacteria associated with Trichodesmium consortia. mSystems. 2023;8:e00742-23.

3. Karl DM. Microbially Mediated Transformations of Phosphorus in the Sea: New Views of an Old Cycle. Annu Rev Mar Sci. 2014;6:279–337.

4. Lanclos VC, Feng X, Cheng C, Yang M, Hider CJ, Coelho JT, et al. New isolates refine the ecophysiology of the Roseobacter CHAB-I-5 lineage. ISME Commun. 2025;ycaf068.

5. Billerbeck S, Wemheuer B, Voget S, Poehlein A, Giebel H-A, Brinkhoff T, et al. Biogeography and environmental genomics of the Roseobacter-affiliated pelagic CHAB-I-5 lineage. Nat Microbiol. 2016;1:16063.

6. Liu Y, Brinkhoff T, Berger M, Poehlein A, Voget S, Paoli L, et al. Metagenome-assembled genomes reveal greatly expanded taxonomic and functional diversification of the abundant marine Roseobacter RCA cluster. Microbiome. 2023;11:265.
